# Neuropeptide Signaling is Required to Implement a Line Attractor Encoding a Persistent Internal Behavioral State

**DOI:** 10.1101/2023.11.01.565073

**Authors:** George Mountoufaris, Aditya Nair, Bin Yang, Dong-Wook Kim, David J. Anderson

**Affiliations:** Division of Biology and Biological Engineering; Program in Computation and Neural Systems; Tianqiao and Chrissy Chen Institute for Neuroscience California Institute of Technology; Howard Hughes Medical Institute Pasadena, CA 91001; Johns Hopkins University School of Medicine; Allen Institute for Brain Science

## Abstract

Internal states drive survival behaviors, but their neural implementation is not well understood. Recently we identified a line attractor in the ventromedial hypothalamus (VMH) that represents an internal state of aggressiveness. Line attractors can be implemented by recurrent connectivity and/or neuromodulatory signaling, but evidence for the latter is scant. Here we show that neuropeptidergic signaling is necessary for line attractor dynamics in this system, using a novel approach that integrates cell type-specific, anatomically restricted CRISPR/Cas9-based gene editing with microendoscopic calcium imaging. Co-disruption of receptors for oxytocin and vasopressin in adult VMH Esr1^+^ neurons that control aggression suppressed attack, reduced persistent neural activity and eliminated line attractor dynamics, while only modestly impacting neural activity and sex- or behavior-tuning. These data identify a requisite role for neuropeptidergic signaling in implementing a behaviorally relevant line attractor. Our approach should facilitate mechanistic studies in neuroscience that bridge different levels of biological function and abstraction.

## Introduction

Innate survival behaviors such as aggression, mating, feeding and defense are driven by internal motivational or emotion states^1–3^, which are experienced in humans as subjective feelings^4,5^. How and where such internal states are encoded in the brain, and how they are causally related to overt behavior, is emerging as a major topic in circuit and systems neuroscience^6,7^.

The study of internal states has been pursued via two general lines of research that have until recently remained relatively separate. One, a “bottom-up” approach, is grounded in perturbational studies and has focused on genetically or pharmacologically based manipulations of genes (e.g., neuromodulators) and neural circuits^6,8,9^ aimed at providing causal explanations for behavioral, psychological or homeostatic internal states^10–12^. The other, a “top-down” approach, is grounded in observational studies, and has defined internal states mathematically through modeling of high-dimensional neural population activity^13,14^. Dynamical system models of low-dimensional neural activity have revealed features such as attractors and integrators that appear to underly many cognitive functions^15–18^. More recently, such models have been applied in behavioral neuroscience as well^19–21^. To test the causal role of such emergent network properties in behavior it is important to understand their neural implementation at the level of specific cell types. This in turn requires integration of these two approaches^22^. Such an integrated approach has thus far been accomplished in very few systems^23,24^.

One important implementation question concerns the mechanism(s) that control slow neural dynamics. Persistent neural activity (on a timescale of seconds to minutes) is a characteristic feature of neural integrators and attractor dynamics^17,25,26^. Two alternative (but not mutually exclusive) implementation mechanisms are typically invoked to explain such persistence: recurrent fast synaptic connectivity or slow neuromodulation^27^. While there is EM connectomic evidence of recurrent connectivity underlying a ring attractor that encodes head direction in *Drosophila*^23,24,28^, to our knowledge there is no evidence of any neuromodulator that controls attractor dynamics in any system.

Neuropeptides comprise a class of evolutionarily conserved^29,30^ neuromodulators that control behavior-specific internal motive states associated with mating^31,32^, aggression^33–35^, social attachment^36^, feeding^37^ and predator defense^38^, as well as other behaviors. Neuropeptides are well known to modulate synaptic strength and neural circuit properties such as patterns of oscillation^39–41^, but their role in implementing persistent activity and attractor dynamics has not been extensively studied in vertebrates. Experiments in *C. elegans* have uncovered the circuit architecture through which neuropeptides control persistent states of locomotor activity^42,43^, but whether they influence population dynamics manifolds identified in that system^44^ is not yet clear. A powerful approach to this question is to integrate specific pharmacological or genetic perturbations of neuromodulatory signaling with simultaneous large-scale recording of neural activity in the same brain region and genetically defined cell type. While this integration has been achieved at the brain-wide scale in *C. elegans*^45^ and larval zebrafish^46^, with few exceptions^47^ it has been difficult to implement in mammalian systems, largely for technical reasons (Supplemental Figure 1A).

Here we describe a novel viral-based strategy that integrates cell type-specific CRISPR/Cas9-based multiplex gene editing^48^ with single unit-resolution calcium imaging in freely behaving adult animals^49^, which we call “CRISPRoscopy”. This method, when combined with dynamical systems modeling, allows investigation of the effects of local inactivation of different neuromodulatory systems on neural population coding, dynamics and behavior, in the same brain region and cell type.

As a proof-of-concept application of this approach, we have examined the role of oxytocin (OXT) and arginine vasopressin (AVP) signaling in a population of ventromedial hypothalamic (VMH) neurons that control aggression^50^. We chose this testbed for several reasons. First, these neuropeptides have been widely implicated in the control of social behaviors^36,51^ (although the role of OXT in aggression has been controversial)^52^. Second, aggression is a robust, naturalistic, evolutionarily conserved and innate social behavior^53^. Third, anatomically restricted^54^ and genetically defined^55,56^ VMH cell populations that play a causal role in aggression have been identified and their activity imaged during aggressive encounters^57–59^. Fourth, VMH neurons are known to express receptors for OXT and AVP^60,61^ and infusion of the latter peptide into VMH can enhance aggression in hamsters^62^. Finally, recent application of dynamical systems modeling^63^ to population recordings from these neurons has revealed an approximate line attractor that represents a scalable and persistent internal state of aggressiveness^20^.

Due to ligand cross-talk^64^ and potential genetic redundancy between *Oxtr* and *Avpr1a*, as a first step to applying this technology we elected to simultaneously edit both genes. We found that genetically disrupting both receptors in VMHvl^Esr1^ neurons^55^ strongly reduced the intensity and frequency of attack behavior, but caused only a modest reduction in overall neuronal activity, and population coding of behavior and intruder sex^59^. In contrast, it strongly reduced persistent neural activity. Dynamical systems modeling revealed that the line attractor was replaced by a slow point corresponding to the animals’ resting state. These data provide the first evidence in any system of a requirement for neuropeptide signaling in the implementation of an attractor.

## Results

### Most Esr1^+^ neurons co-express Oxytocin and Vasopressin receptors and respond to these peptides *ex vivo*

The expression of Oxtr and Avpr1a has been extensively mapped in the brains of multiple species by radioligand binding, in situ hybridization and knock-in reporters^60,61,65–68^. To determine whether these receptors are co-expressed in individual VMH neurons we examined a single cell RNAseq (scRNAseq)-based atlas of VMHvl transcriptomic cell types anatomically validated by smFISH^69^. We observed co-expression of *Oxtr* and *Avpr1α* mRNA transcripts (but not of *Avpr1β* or *Avp2r* transcripts) within individual neurons belonging to several Esr1^+^ transcriptomic clusters (Figure 1B and Supplemental Figure S1B, E). All Esr1^+^ VMHvl clusters contained cells expressing *Oxtr*, with most clusters (5/8) containing cells that co-expressed *Avpr1a* transcripts (Figure 1C-D and Supplemental Figure S1C). Overall, ∼78% of Esr1^+^ cells in VMHvl expressed Oxtr mRNA, and ∼61% of these cells also expressed Avpr1a mRNAs (Figure 1E). However, none of the VMHvl clusters expressed *Oxt* or *Avp* transcripts, indicating that the source(s) of the peptides must be extrinsic to the nucleus (Supplemental Figure S1D).

**Figure 1.**
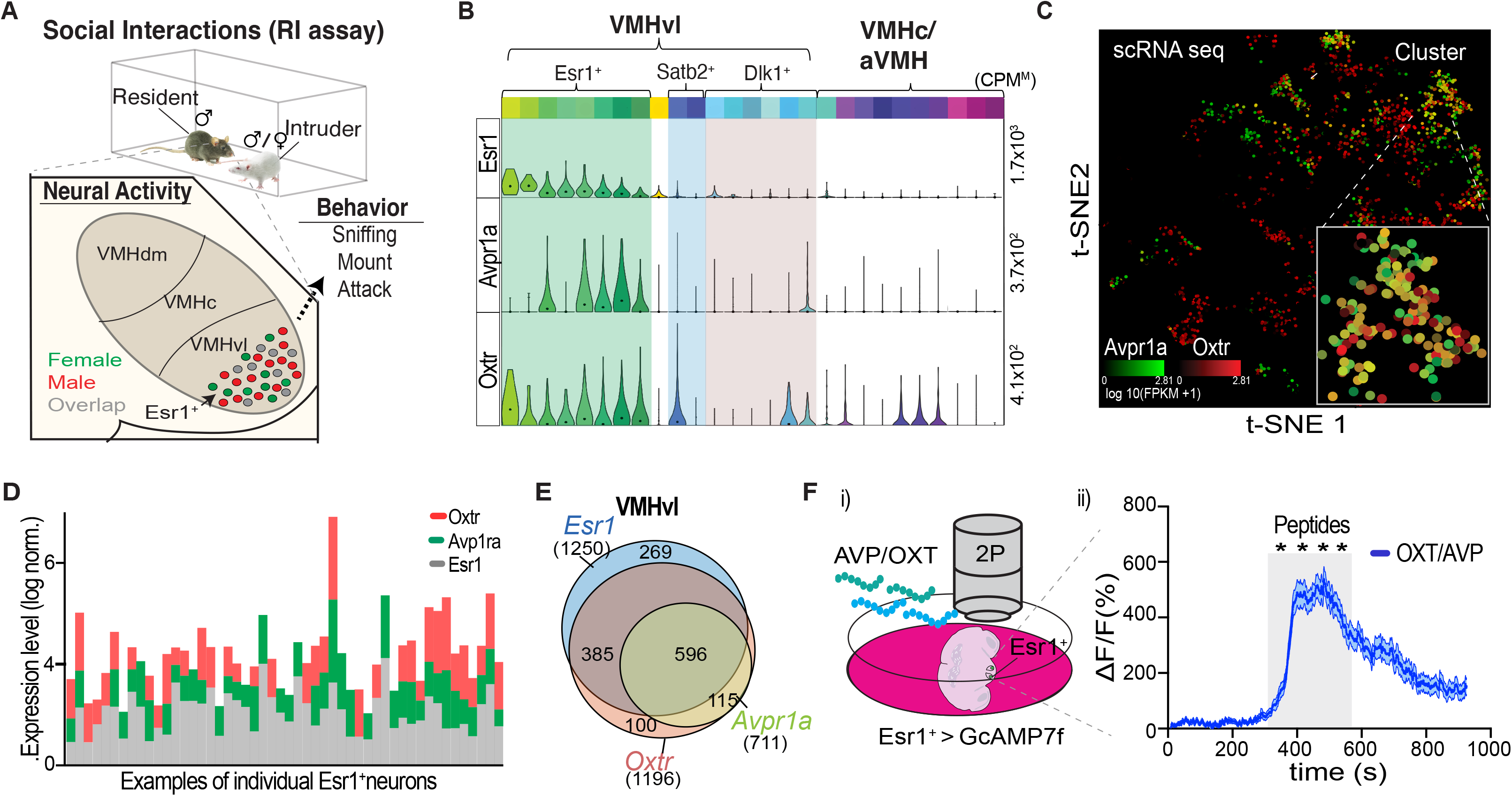
VMHvl^Esr1^ neurons co-express Oxytocin and Vasopressin receptors and respond to these peptides *ex vivo*. **A)** A graphic illustration of the resident intruder behavioral assay used in this study (top). A graphic illustration of male-(red), female-(green), and mixed-selective (grey) VMHvl^Esr1^ responses in male residents during interactions with male vs. female intruders. Optogenetic interrogation of VMHvl^Esr1^ cells in male mice controls appetitive (sniffing) and consummatory (mount and attack) behaviors towards conspecific intruders. **B)** Violin plots illustrating the expression of *Esr1, Avpr1a* and *Oxtr* mRNAs in single cell gene clusters (top-colored boxes); “max CPM”, maximum counts per million reads. Green, blue, and red boxes highlight the expression of *Esr1, Avp1ra*, and *Oxtr* in the three major VMHvl gene clusters. **C)** t-SNE plots illustrating the distribution of *avpr1a* and *oxtr* mRNAs (ii) in single cells from the VMH. All cells visible in the inset are Esr1^+^ (see Supplemental Figure S1C). **D)** Example of ∼50 individual VMH cells co-expressing *Esr1, Avpr1a*, and *Oxtr* mRNAs. **E)** Venn diagram of *esr1, avpr1a*, and *oxtr* mRNA expressing neurons within the VMHvl. The size of each circle is proportional to the number of neurons. Numbers represent individual neurons expressing each receptor gene. Analyses in (B-E) are based on data originally reported in Kim et al. (2019). **F)** Graphic illustration of the two-photon *ex vivo* calcium imaging experiments. (i) Acute brain slices from male ESR1-2A-CRE animals that express Cre-dependent GCaMP7f virus were perfused with OXT and/or AVP neuropeptides. (ii) Quantification of GCaMP7f calcium responses in Esr1^+^ cells to a 400nM cocktail of AVP/OXT (n=124 cells, 4=mice). Gray rectangle depicts the duration of the bath application of peptides. For responses to individual peptides see Supplemental Figure S1F. Statistics: Values plotted as mean ± S.E.M. ****p<0.0001 Mann-Whitney test

To explore the effect of OXT and AVP on the physiological activity of VMHvl neurons, we used a system for 2-photon imaging system of acute VMH slice preparations^55^ (Figure 1Fi). This system enabled us to record Ca^2+^ traces in brain slices from ESR1-2A-CRE animals^55^ expressing a CRE-dependent Ca^2+^ indicator (GCaMP7f)^70^. Perfusing a mixture of 400nM (each) OXT plus AVP or individually administering these peptides elicited strong (ΔF/F ∼500-600%) responses in VMHvl^Esr1^ cells (Figure 1Fii and Supplemental Figure S1F-G). These findings demonstrated the presence of functional AVP and OXT receptors in VMHvl^Esr1^ neurons, consistent with prior electrophysiological studies in mice and guinea pigs^71,72^. Furthermore, it established a system for evaluating cellular responses to OXT and AVP following CRISPR/Cas9 based mutations of their receptors.

### Oxtr/Avpr1a-mediated neuropeptidergic signaling in VMH is required for male territorial aggression

The likelihood of functional redundancy or developmental compensation between OXTR and AVPR1a in mediating responses to OXT and/or AVP^64,71^ prompted us to design a multiplex viral CRISPR/Cas9 approach to concurrently target both receptors in the same cells. To do this, we modified previously described vectors^73^ to express two different gRNAs targeting *Oxtr* and *Avpr1a* from four distinct Pol III promoters in either a lentiviral (LV) or an AAV vector (Figure 2A; see Methods). We used two different gRNAs for each receptor gene since this approach has been reported to significantly increase *in vivo* gene editing efficiency in the mammalian nervous system^74^. Each gRNA was designed to target either the *Oxtr* or the *Avpr1a* coding region, and this specificity was confirmed using the T7 endonuclease assay^75^ (Supplemental Figure S2A).

**Figure 2.**
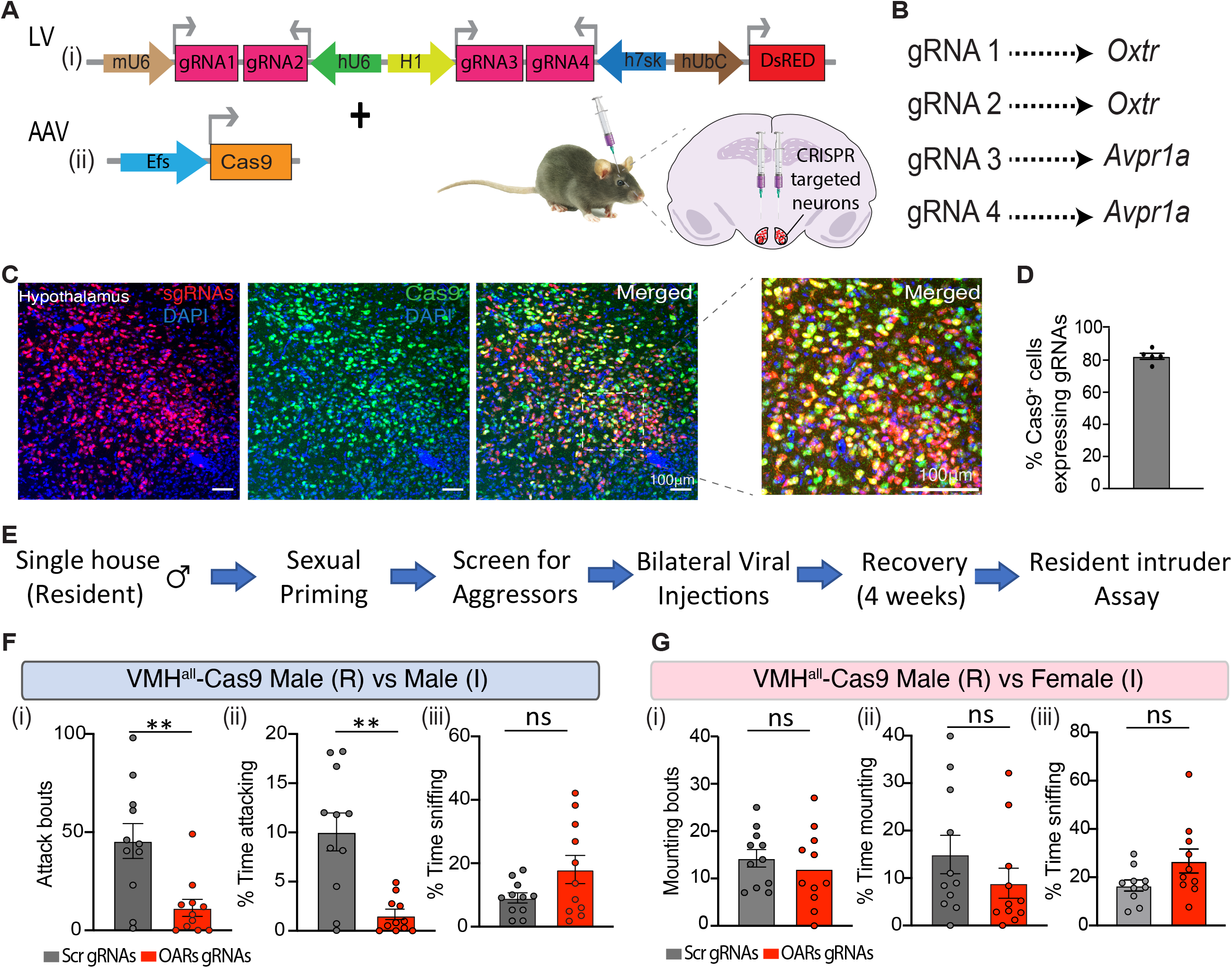
CRISPR/Cas9 based co-perturbation of *Oxtr* and *Avpr1a* reduces territorial aggression in males. **A)** A schematic of the strategy for region-restricted multiplex CRISPR/Cas9 gene editing *in vivo*. A mixture of the two viruses was used-A Lentivirus (LV) constitutively expressing four different gRNAs and the red fluorescence marker DsRed (i). A Adenovirus (AAV) constitutively expressing the Cas9 protein (See Methods) (ii). Male residents were injected bilaterally in the VMH with the OARs-Cas9 (experimental group) or Scr-Cas9 (control group) viral mixture. **B)** Two different gRNAs target each GPCR receptor gene. Scrambled gRNA sequences were used as negative controls. **C)** Immunostaining against the DsRED (red) and Cas9 (green) proteins in coronal hypothalamic sections from animals co-injected with LV gRNA virus and the Cas9 AAV. Sections are counterstained with Dapi (blue). **D)** Quantification of Cas9+ cells (Aii) infected with the gRNA (Ai) virus in the hypothalamus. n= 3 mice. **E)** Diagram of the behavioral paradigm used for testing the effects of Oxtr and Avpr1a mediated signaling in social behaviors. **F)** Quantification of the number (i) and the total time spent attacking (ii), or sniffing (iii) male intruders in experimental and control mice. **G)** Quantification of the number (i) and the total time spent mounting (ii) or sniffing (iii) female intruders in experimental and control mice. n= 11 mice per group. Statistics: Mann-Whitney test was performed **p≤0.01.

While OXT, AVP and their receptors have been studied extensively in rodent aggression using pharmacologic reagents and organismal gene knockouts,^31,52,76–79^ there are no reports of a specific requirement in offensive aggression for either *Oxtr* and/or *Avpr1a* in murine VMH. Because pharmacological blockers are difficult to restrict to small anatomical regions due to diffusion that cannot be visualized following injection, we initially sought to use CRISPR/Cas9-mediated gene editing to evaluate whether these receptors are functionally relevant to this behavior in VMH *in vivo*. Generating viruses with the high titers necessary for *in vivo* applications (see Methods) precluded the inclusion of a Cas9 cDNA in the same vector that encoded two different gRNAs each against *Oxtr* and *Avpr1a*. Therefore, Cas9 was delivered by co-injection of a separate AAV (Figure 2Aii). To target VMHvl neurons broadly for this initial experiment, we employed a constitutive version of the system. We used a LV-based vector to express the 4 different gRNAs, because we found that its anatomical spread is more restricted than that of AAVs.

Examination of hypothalamic sections from animals co-injected with the gRNA-DsRed LV and the Cas9 AAV using DsRed fluorescence and anti-Cas9 immunostaining indicated that ∼80% of Cas9^+^ cells were co-infected with the gRNA virus (Figure 2C, D). Assessing the efficiency of co-disrupting both receptors *in vivo* at the protein level was not feasible due to a lack of equally specific immune reagents for both OXTR and AVPR1a. Furthermore, because the small 1-2 nucleotide DNA insertions or deletions (INDELs) created by CRISPR/Cas9 editing do not necessarily result in reduced levels of mRNA expression from the targeted genes^80^, assessing editing by *in situ* hybridization is not a reliable indicator of targeting efficiency. As described in the next section, we tested the functional efficacy of *Oxtr/Avpr1a* co-editing via calcium imaging in acute VMH brain slices, and indeed found a strong suppression of physiological responses to OXT+AVP (Supplementary Figure S3B and Figure 3A).

**Figure 3.**
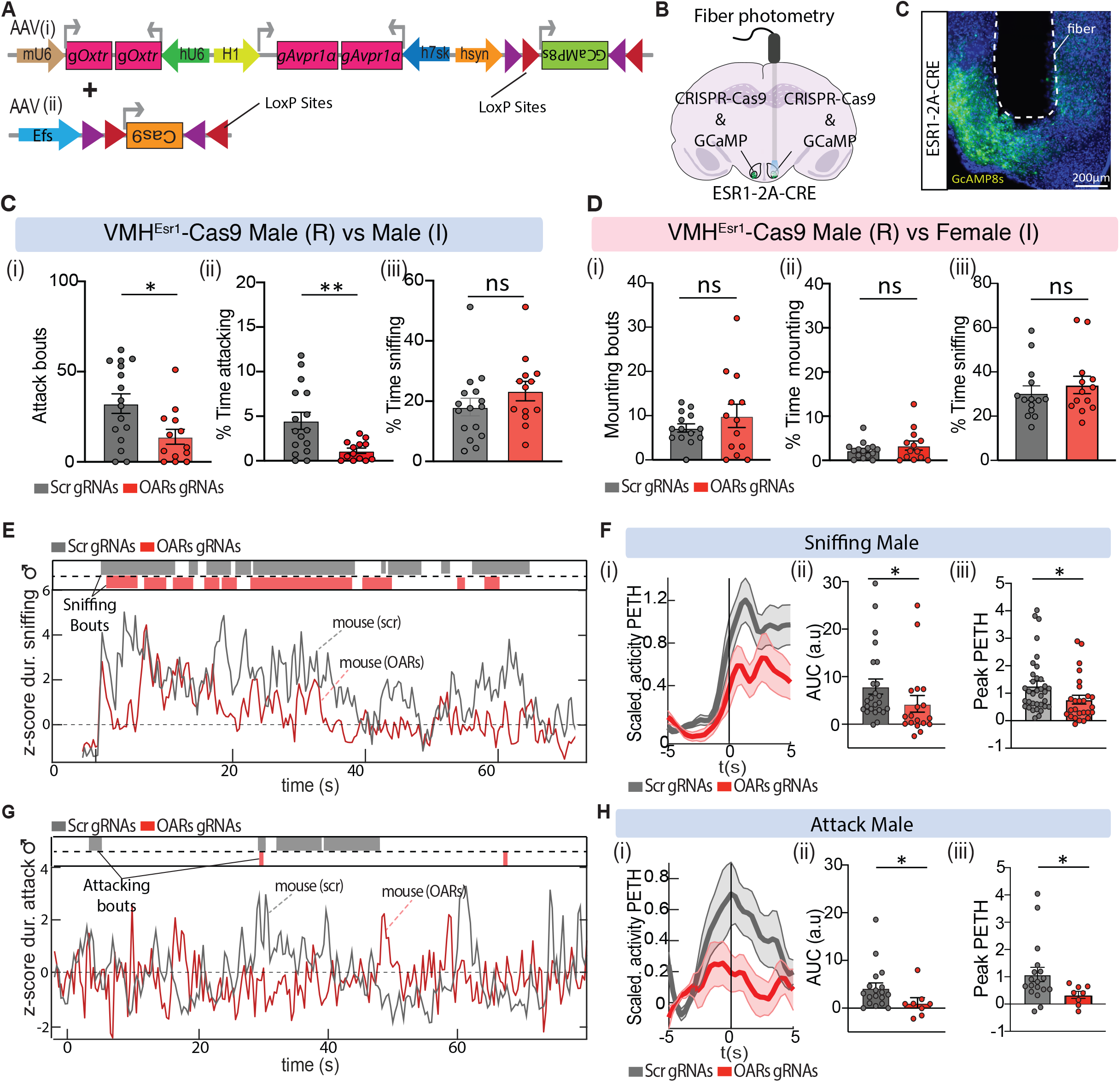
Co-perturbation of OXT/AVP signaling alters VMHvl^Esr1^ bulk calcium activity. **A)** A schematic of the strategy for cell type specific Cre-dependent multiplex CRISPR/Cas9 gene editing and calcium imaging *in vivo*. As before, two different gRNAs target each GPCR receptor gene. Scrambled gRNA sequences were used as negative controls. Cre-dependent expression of GCaMP8s (i) and Cas9 proteins (See Methods) (ii). **B)** Graphic illustration of unilateral fiber photometry recording experiments in ESR1-2A-CRE male residents that are bilaterally co-injected with Cre-dependent Cas9 AAV and a Cre-dependent OARs-GCaMP8s gRNA (experimental group) or Scr-GCaMP8s gRNA AAVs (control group) in the VMHvl. Coronal histology section showing the expression of GCaMP8s (green) in VMHvl. Section counterstained with Dapi (blue). **C)** Quantification of the number (i) and the total time spent attacking (ii) or sniffing (iii) male intruders in experimental and control mice. n=16 control and n=13 experimental mice. **D)** Quantification of the number (i) and the total time spent mounting (ii) or sniffing (iii) female intruders in experimental and control mice. n=16 control and n=13 experimental mice. **E)** Examples of fiber photometry traces from a control and experimental mouse during male-directed sniffing. **F)** Z-scored behavioral triggered average (BTA) of VMHvl^Esr1^ activity (i), the average area under the curve (AUC) (ii) and the peak PETH signal during male-directed sniffing (1 min). Each dot represents a bout. n=5 mice per group. **G)** Examples of fiber photometry traces from a control and experimental mouse during attack. **H)** Z-scored BTA of VMHvl^Esr1^ activity (i), the average area under the curve (AUC) (ii) and peak PETH signal during attack. n=5 per group. Statistics: Values plotted as mean ± S.E.M. Mann-Whitney test was performed *p≤0.05 **p≤0.01;

Next, we examined social behaviors in mice injected in VMH bilaterally with the OAR-gRNA LV and a Cas9 AAV, using a standard resident intruder (RI) assay (Figure 1A). We used single-housed, sexually experienced wild-type C57BL/6N resident males pre-selected for adequate aggressiveness (Figure 2E, see Methods). Control animals were co-injected bilaterally with the scrambled gRNA (Scr gRNA) and Cas9 viruses. Experimental mice displayed a notable reduction in aggression towards male intruders, as evidenced by significant decreases in the number and time-varying probability of attack bouts, the total time spent attacking and the average duration of each attack bout; in addition, the latency to the 1st attack bout was significantly increased (Figure 2Fi-ii and Supplemental Figure S2B, Ci, ii). These behavioral effects were not due to defects in locomotor activity since average velocity during attack episodes was similar between experimental and control mice (Supplemental Figure S2Ciii). In contrast, the time spent in close investigation (sniffing) of male intruders did not differ significantly between experimental and control male residents (Figure 2Fiii). Experimental males also did not differ significantly from controls in their sniffing or mounting behavior towards female intruders (Figure 2G and Supplemental Figure S2D).

These data suggest that *Oxtr* and/or *Avpr1a* expressed in VMHvl neurons play a requisite and selective role in aggressive behavior. Furthermore, they motivated us to analyze next how disrupting these receptors specifically in Esr1 neurons affects behavior, neural activity, population coding and network dynamics *in vivo*.

### Altered VMHvl^Esr1^ neural activity during social interactions in mice with perturbed OXT/AVP signaling

Male VMHvl^Esr1^ neural activity normally increases during sniffing and attack towards an intruder male^57,59,81^. We therefore sought to analyze the activity of these neurons via calcium imaging in mice with or without co-disruption of *Oxtr* and *Avpr1a*. To restrict CRISPR-based gene editing to the same cell population that we wished to image, we constructed an AAV vector that encodes both the gRNAs and a Cre-dependent GCaMP8s (Figure 3A, upper). In mice co-injected with a Cre-dependent Cas9 AAV, cells co-infected with both viruses are expected to undergo gene editing. The use of Cre-dependent AAVs in these experiments yields higher levels of expression than LVs as well as cell type specificity. Double-labeling with antibodies to GFP (GCaMP8s) and Cas9 in mice co-injected with the two Cre-dependent AAVs (Figure 3A) indicated that ∼66% of GCaMP8s-expressing Esr1 neurons were Cas9^+^ (Supplementary Figure S3Ai-ii).

As mentioned above, *in vivo* quantification of the frequency and efficiency of *Oxtr/Avpr1a* gene disruption on a cell-by-cell basis (either by immune-labeling *or in situ* hybridization) was precluded for technical reasons. As an alternative approach to verify the disruption of normal signaling responses to OXT and AVP in Esr1 neurons with co-targeting of *Oxtr* and *Avpr1a*, we performed *ex vivo* calcium imaging of VMH slices (Figure 1Fi) from ESR1-2A-CRE mice co-injected with the Cas9 and Cre-dependent gRNAs-GCaMP8s viruses (Figure 3Ai). Physiological responses to the application of mixed AVP and OXT peptides were significantly attenuated (∼2.5-fold) in slices from experimental mice expressing *Oxtr/Avpr1a* gRNAs in Esr1^+^ cells, compared to slices from control mice expressing a scrambled (Scr) gRNA in these cells (Supplementary Figure S3B). Thus, co-disruption of *Oxtr/Avpr1a* using our dual viral-based CRISPR/Cas9 system can reduce physiological responses to OXT and AVP in VMHvl^Esr1^ neurons, although they do not eliminate such responses completely.

We next used the Cre-dependent gene editing and GCaMP imaging system to test whether *Oxtr/Avpr1a* mediated signaling is required selectively in Esr1 neurons during aggression, and to examine simultaneously how neural activity in these cells is affected. Similar to the results obtained initially with the Cre-independent system, bilateral co-disruption of *Oxtr/Avpr1a* in VMHvl^Esr1^ neurons significantly reduced most metrics of aggression towards male intruder compared to control mice, although to a somewhat lower extent (Figure 3Ci, ii; Supplementary Figure 3C). Using fiber photometry to measure bulk calcium signals (Figure 3B, C), we observed a significant reduction in overall VMHvl^Esr1^ neuronal activity during the infrequent attack bouts exhibited by experimental mice (Figure 3G, compare red vs. grey rasters) (Figure 3H). A reduction in VMHvl^Esr1^ activity during sniffing episodes was also observed in these mice (Figure 3E-F), even though the duration of investigative behavior remained unaffected (Figure 3Ciii). In contrast, during male-female interactions, calcium signals in experimental males were not significantly different from controls (Supplementary Figure S3E-F), consistent with the lack of an effect on mounting behavior (Figure 3D; Supplementary Figure S3D). Thus, males with bilateral co-disruption of Oxtr/Avpr1a-mediated signaling in VMHvl^Esr1^ neurons exhibited a reduction in both aggression and neural activity. However, these data did not distinguish whether the reduced activity was a cause or a consequence of the diminished aggressive behavior.

### Single cell “CRISPRscope” imaging of VMHvl^Esr1^ neurons with co-disruption of *Oxtr/Avpr1a*

To investigate how co-editing of *Oxtr/Avpr1a* affects activity in individual VMHvl^Esr1^ neurons, we imaged Ca^2+^ activity using a miniature head-mounted microscope^49^ in Esr1-2A-CRE males co-injected with the experimental or control virus pairs. We call this approach “CRISPRoscopy.” Because there are virtually no commissural connections between VMHvl^58^, unilateral loss-of-function manipulations of this nucleus are typically compensated by the unmanipulated side. To avoid any influence on neural activity of reduced aggressive behavior caused by *Oxtr/Avpr1a* co-disruption, we performed calcium imaging and functional perturbations unilaterally in freely behaving animals (Figure 4A), a previously validated strategy^58^.

**Figure 4.**
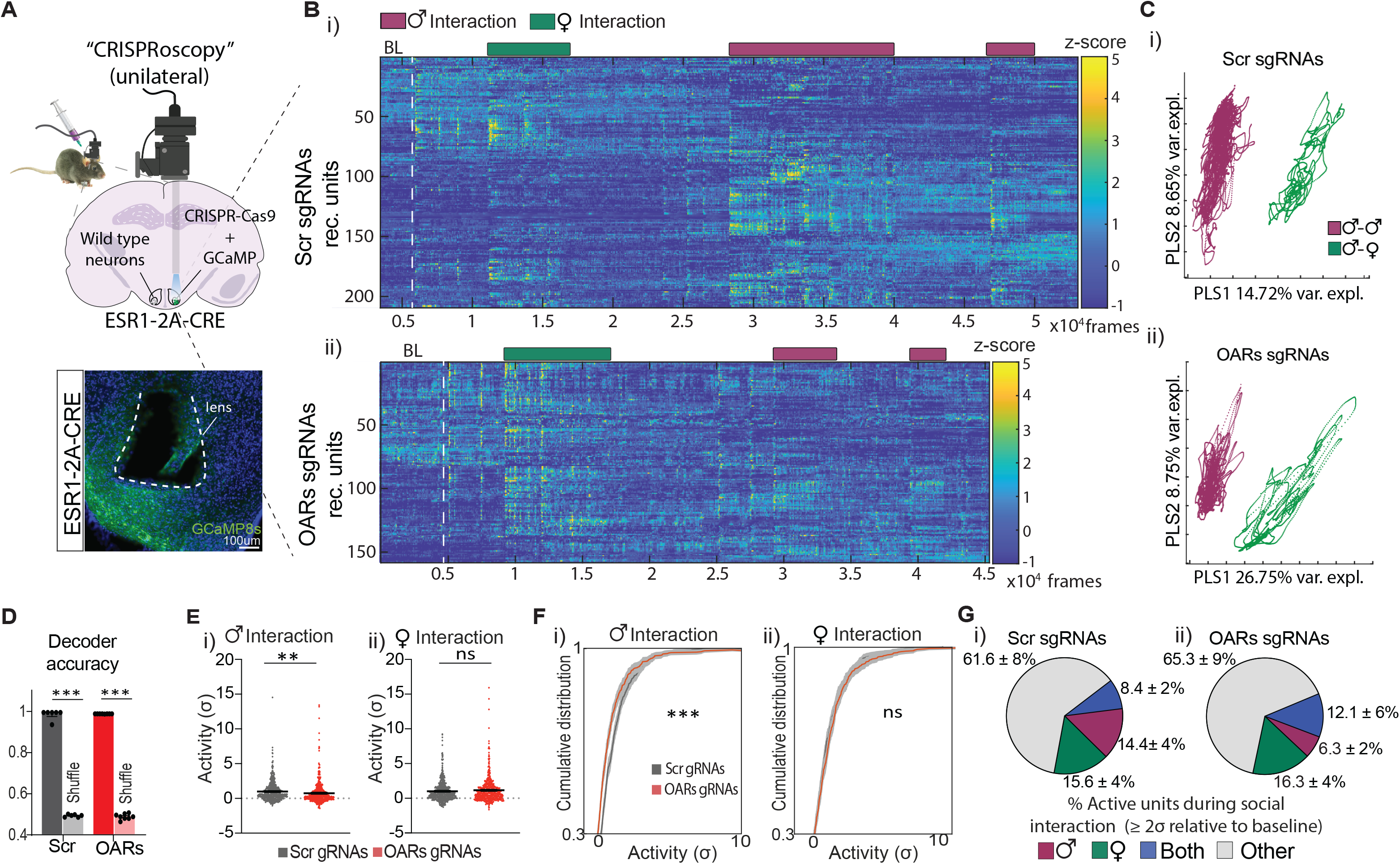
Single cell “CRISPRscope” imaging of VMHvl^Esr1^ neurons with co-disruption of *Oxtr/Avpr1a*. **A)** A graphic illustration of “CRISPRoscopy.” Male ESR1-2A-CRE residents unilaterally co-injected with a Cre-dependent Cas9 AAV and Cre-dependent OARs-GCaMP8s AAV (experimental group) or Cre-dependent Scr-GCaMP8s AAV (control group) in the VMHvl (top). Coronal histology section showing the expression of GCaMP8s (green) in VMHvl (bottom). Section counterstained with Dapi (blue). **B**) Example of micro-endoscope single unit z-scored responses towards female or male intruders from control (i) and experimental (ii) male residents. **C)** Example of VMHvl^Esr1^ ensemble representations of intruder sex, for a control (i) and experimental (ii) male, projected onto the first two axes of a PLS regression against intruder sex. Traces are colored by intruder sex identity. The percentage of variance explained by the first two PLS components is noted for each mouse. **D**) Accuracy of frame-wise decoders predicting the sex of the intruder trained on VMHvl^Esr1^neural activity in control and experimental animals. **E**) Average single VMHvl^Esr1^unit (σ) responses and **F)** cumulative distribution of VMHvl^Esr1^ activity (σ) relative to pre-intruder baseline, towards male (i) or female (ii) intruders in control and experimental mice during 1 minute of interaction. **G)** Percentage of male- or female selective, - or co-active VMHvl^Esr1^ units (≥ 2σ above the pre-intruder baseline) in control and experimental mice. n=5 control, n=7 Oxtr/Avp1ra targeted animals. Statistics: Values plotted as means with ± S.E.M in (F) and in (G). Nested Mann-Whitney test was performed except (F), where nested Kolmogorov–Smirnov test was used. **p≤0.01 ***p≤0.001 ****p≤0.0001

### Effect of Oxtr/Avpr1a co-editing on intruder sex-specific representations during social encounters

Our previous single unit calcium imaging studies have shown that in socially experienced males VMHvl contains distinct Esr1^+^ neural ensembles activated in the presence of males vs. females, with some units showing mixed sex-selectivity ^2,57–59^. This separation was also clear in raster plots of VMHvl^Esr1^ units imaged in control vs experimental males (Figure 4B). To quantify the proportion of sex-preferring neurons, we measured unit activity during the first two minutes after the introduction of a male or female intruder in two ways: either by z-scoring (relative to the cell’s mean fluorescence over the entire recording period), or by the change in fluorescence relative to the mean pre-intruder baseline ^57,58^ (in units of σ; see Methods).

During interactions with a male intruder, the experimental cumulative distribution function (ECDF) and mean activity of all units (pooled from n=4 control and n=7 experimental animals) were slightly but significantly decreased in experimental mice (Figure 4Ei, Fi; Supplementary Figure 4Ci). However, the mean activity among cells considered as “active” (≥ 2σ above baseline^59^) did not differ between control and experimental mice (Supplemental Figure S4D). During interactions with females there was no significant difference in activity (measured in σ above baseline) between experimental and control animals (Figure 4Eii, Fii), although z-scored activity showed a slight but significant increase (Supplementary Figure 4Cii).

Next, we measured the percentage of male-selective (activity ≥ 2σ during male but not female interactions) and mixed selectivity (activity ≥ 2σ during both male and female interactions) neurons within the Esr1^+^ population during male-male interactions. The combined percentage of all male-activated neurons was slightly smaller in experimental than control mice (34.6% vs. 38.3%, respectively; Figure 4G and Supplemental Figure S4E), while there was a ∼56% reduction in the fraction of male-selective neurons (6.3 ± 2% vs. 14.4 ± 4%, respectively; Figure 4G). Conversely, during female interactions the fraction of female-selective and mixed selectivity neurons was increased by ∼4% and ∼44%, respectively, in experimental mice (Figure 4G and Supplemental Figure S4E). Together these data are suggestive of a shift in sex-specific tuning from male-selective to mixed selectivity, a conclusion consistent with choice probability analysis (see below).

To determine whether this shift affected the ability to accurately decode intruder sex at the population level, we performed dimensionality reduction using partial least-squares (PLS) regression. This analysis revealed a clear separation of responses during encounters with males vs. females in both control and experimental animals (Figure 4C; Supplemental Figure S4A-B). In addition, linear SVM decoders trained on imaging data from either control or experimental mice correctly predicted intruder sex with virtually 100% accuracy (Figure 4D). Thus in socially experienced animals, co-targeting of *Oxtr*/*Avpr1a* in VMHvl^Esr1^ cells does not disrupt the population coding of intruder sex^59^, despite the altered sex-selectivity of some of these units.

### Social behavior-selective tuning of VMHvl^Esr1^ neurons with co-disruption of *Oxtr*/*Avpr1a*

Next, we examined the effect of co-disruption of *Oxtr/Avpr1a* on VMHvl^Esr1^ neuronal activity during the different behavioral phases of social interactions with males or females: appetitive (sniffing) or consummatory (attack or mounting, respectively). The ECDF and average single unit activity during male-directed sniffing or attack was slightly but significantly lower in experimental than in control mice (Figure 5Ai, ii, Ci, ii and Supplementary Figure S5Ai, Bi-ii). In contrast, during social interactions with females’ activity during sniffing and mounting was similar (Figure 5Aiii, iv, Ciii, iv and Supplementary Figure S5Biii-iv). As an additional approach, we quantified the average activity in peri-event time histograms (PETHs) for each type of behavior after subtracting the pre-behavior baseline, using z-scored data (see Methods). The mean activity during male-directed sniffing or attack was slightly but significantly lower in experimental than in control mice (Supplemental Figure S5Ci, ii), while it was significantly higher during female-directed sniffing, but unchanged during mounting (Supplemental Figure S5Ciii, iv). In summary, the activity of VMHvl^Esr1^ units during different social behaviors was either unchanged or only slightly different between experimental and control animals, with minor decreases or increases during male-vs. female-directed behaviors, respectively.

**Figure 5.**
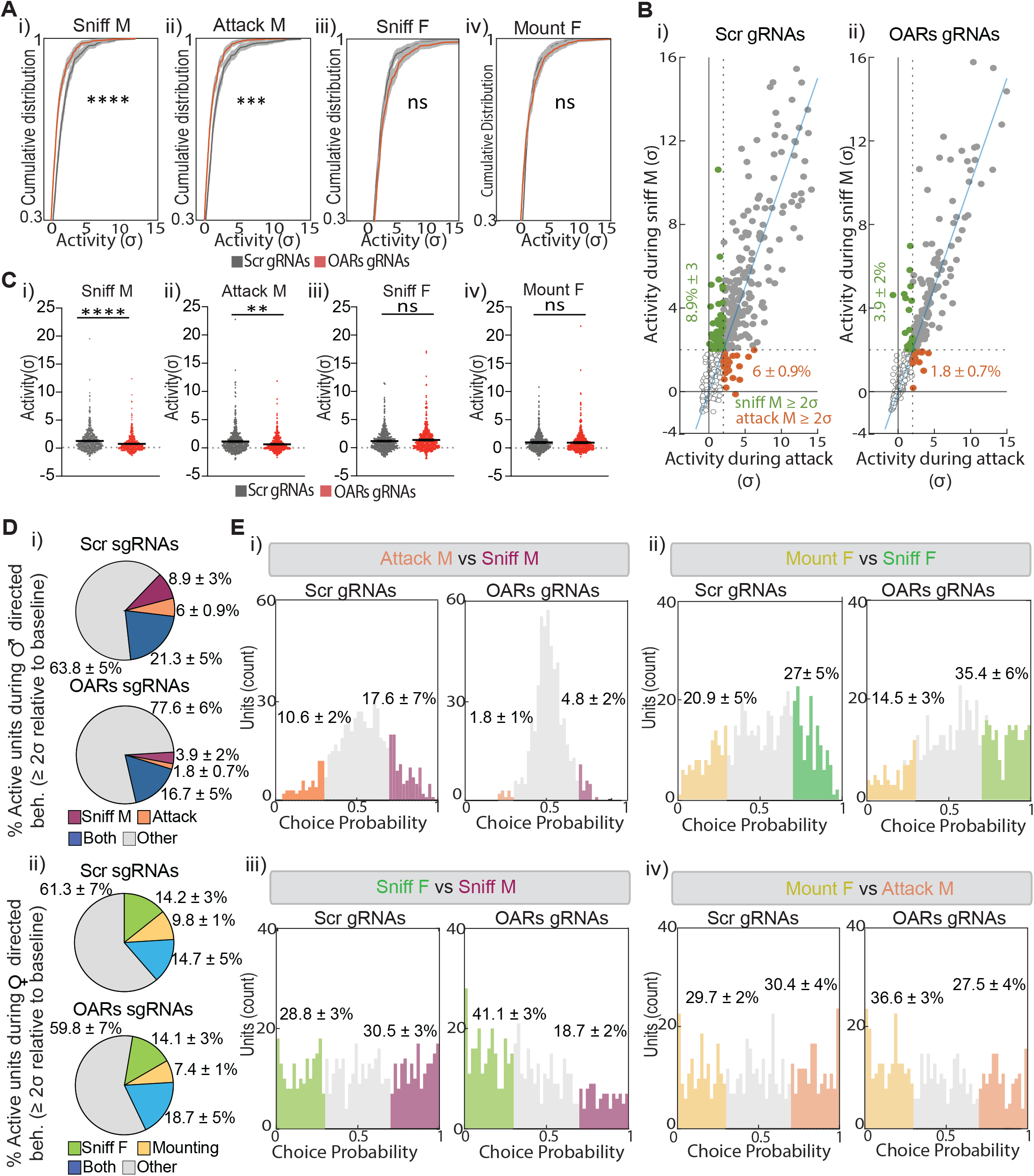
Social behavior-selective activity and tuning of VMHvl^Esr1^ neurons with co-disruption of *Oxtr*/*Avpr1a*. **A)** Cumulative distribution of VMHvlEsr1activity (σ) during male- (sniffing and attack) or female (sniffing and mounting) -directed behaviors in control and experimental mice relative to the pre-intruder baseline. **B**) Scatter plots of single VMHvl^Esr1^ unit activity (σ) during male-directed sniffing or attack in control and experimental mice. Green data points depict units with ≥ 2*σ* activity during male-directed sniffing and < 2*σ* activity during attack, relative to the pre intruder baseline activity. Red data points depict units with ≥ 2 activity during attack and < 2*σ* activity male-directed sniffing, relative to the pre intruder baseline activity. **C)** Average activity (σ) of single VMHvl^Esr1^ units during male-(sniffing and attack) or female (sniffing and mounting)-directed behaviors in control and experimental mice relative to the pre-intruder baseline. **D**) Percentage of VMHvl^Esr1^ units active (defined as ≥ 2*σ* relative to pre intruder baseline) during male-(i) or female-directed behaviors (ii) in control and experimental mice. **E**) Choice probability histograms of VMHvl^Esr1^ tuning during male- and female-directed behaviors in control and experimental mice. Statistics: Values plotted as means in (A) (± S.E.M), (D) and (E). Nested Kolmogorov–Smirnov test was used in (A), whereas the Nested Mann-Whitney test was performed in (C). **p≤0.01 ***p≤0.001 ****p≤0.0001

We next examined the proportion of behavior-selective units (defined as units with activity ≥ 2σ above pre-intruder baseline during, e.g., sniff but not attack or vice-versa)^57–59^. As we previously showed, a relatively small fraction of VMHvl^Esr1^ neurons was selective for sniff or attack (∼2.5-10%), with the majority showing mixed behavioral selectivity (Figure 5Bi)^57–59^. In experimental mice, during male interactions the fraction of sniff-selective units was reduced by ∼40% relative to controls (3.9 ± 2% in OARs gRNAs mice vs. 8.9 ± 3% in Scr gRNAs mice) while the proportion of attack-selective units was reduced by ∼70% (1.8 ± 0.7% vs. 6 ± 0.9%; Figure 5B, Di). The fraction of neurons exhibiting mixed selectivity (i.e., active during both behaviors) was moderately reduced (∼22%; Figure 5Di). Overall, there was an ∼38% reduction in the fraction of active (≥ 2σ above baseline) units during all male-directed behaviors (from 36.2% in control to 22.4% in experimental mice). In contrast, the fraction of female-directed sniffing- or mount-selective units was similar between control and experimental males (Figure 5Dii).

As an alternative metric of a cell’s behavioral tuning, we calculated its choice probability (CP): a cell was considered “tuned” to one of two pairwise-compared behaviors if it exhibited a CP > 0.7 that was significantly different (p < 0.5) from shuffled data^59^. We observed an ∼80% and ∼70% reduction in the percentage of attack- and sniff-tuned units, respectively, in experimental vs. control mice during male-male interactions (Figure 5Ei), with a concomitant ∼18% increase in the fraction of units exhibiting mixed selectivity (CP for sniff vs. attack ≤ 0.7; Figure 5Ei, gray bars and Supplementary Figure S5D). In contrast, the percentage of units tuned to sniffing females (vs. sniffing males) was increased in experimental mice (28.8 ± 3% vs. 41.1 ± 3%; Figure 5Eiii). We also observed a slight increase in the fraction of sniff female (vs. mount female) -tuned units and mount female (vs. attack male) -tuned units in experimental vs. control animals (Figure 5Eii). This analysis suggests that perturbation of normal OXTR/AVPR1a-mediated signaling decreases the relative number of units tuned to specific male-directed behaviors and increases the fraction of cells tuned for female-directed behaviors.

Despite these quantitative changes in behavior-selective tuning, there was no significant difference between control and experimental mice in the performance of linear decoders trained to identify and distinguish attack from sniffing behavior based on the activity of VMHvl^Esr1^ units (Supplemental Figure S5E). Decoders trained on data from each group were able to accurately distinguish these behaviors in held out testing data with close to 100% accuracy. Thus, the population coding of social behavior by VMHvl^Esr1^ neurons^59^ is not affected by co-disruption of *Oxtr/Avpr1a*.

### VMHvl^Esr1^ line attractor dynamics require Oxtr/Avpr1a-mediated signaling

In addition to the level of activity and feature-specific tuning, neural dynamics can play an important role in the neural coding of cognitive function or internal state^17^. Using unsupervised linear dynamical systems modeling^63^, we recently discovered an approximate line attractor in VMHvl neural state space that encodes a low-dimensional, scalable representation of aggressiveness ^20^. This line attractor is implemented by a subset of VMHvl^Esr1^ neurons (∼20-25%) whose collective activity ramps up as social interactions escalate to attack and thereafter decays with a long (∼100s) time constant^20^. This long time constant reflects persistent activity in this subset. Since neuromodulatory signaling has been implicated in some forms of persistent neural activity^27,82,83^, we investigated whether line attractor dynamics during aggression are altered when OXTR/AVPR1a-mediated signaling is perturbed.

We fit a recurrent switching linear dynamic system (rSLDs) model^84^ using data from animals with at least 31 imaged units, thereby reducing the dimensionality of the data to 5 latent factors and three states (S1-S3) which captured ∼85% of the observed variance in neural activity. Attack behavior occurred during a single rSLDs state in both control and experimental males. In control mice, the time constant (τ) of the 1^st^ rSLDS dimension (derived from the first Eigenvalue of the fit dynamics matrix; see Methods) was much higher than that of the 2^nd^ dimension (∼100-120s vs. ∼40s; Figure 6A). This yielded a line attractor score (calculated as the log_2_ of the ratio of the τ ‘s of the 1^st^ and 2^nd^ dimensions) of ∼1.5 (Figure 6F), similar to that observed in mice without CRISPR/Cas9 gene editing^20^. In contrast, in experimental mice the first two rSLDS dimensions had statistically indistinguishable τ values (<50s; Figure 6C), due to a reduced 1^st^ dimension τ (Supplementary Figure 6D) and consequently a line attractor score close to zero (Figure 6F).

**Figure 6.**
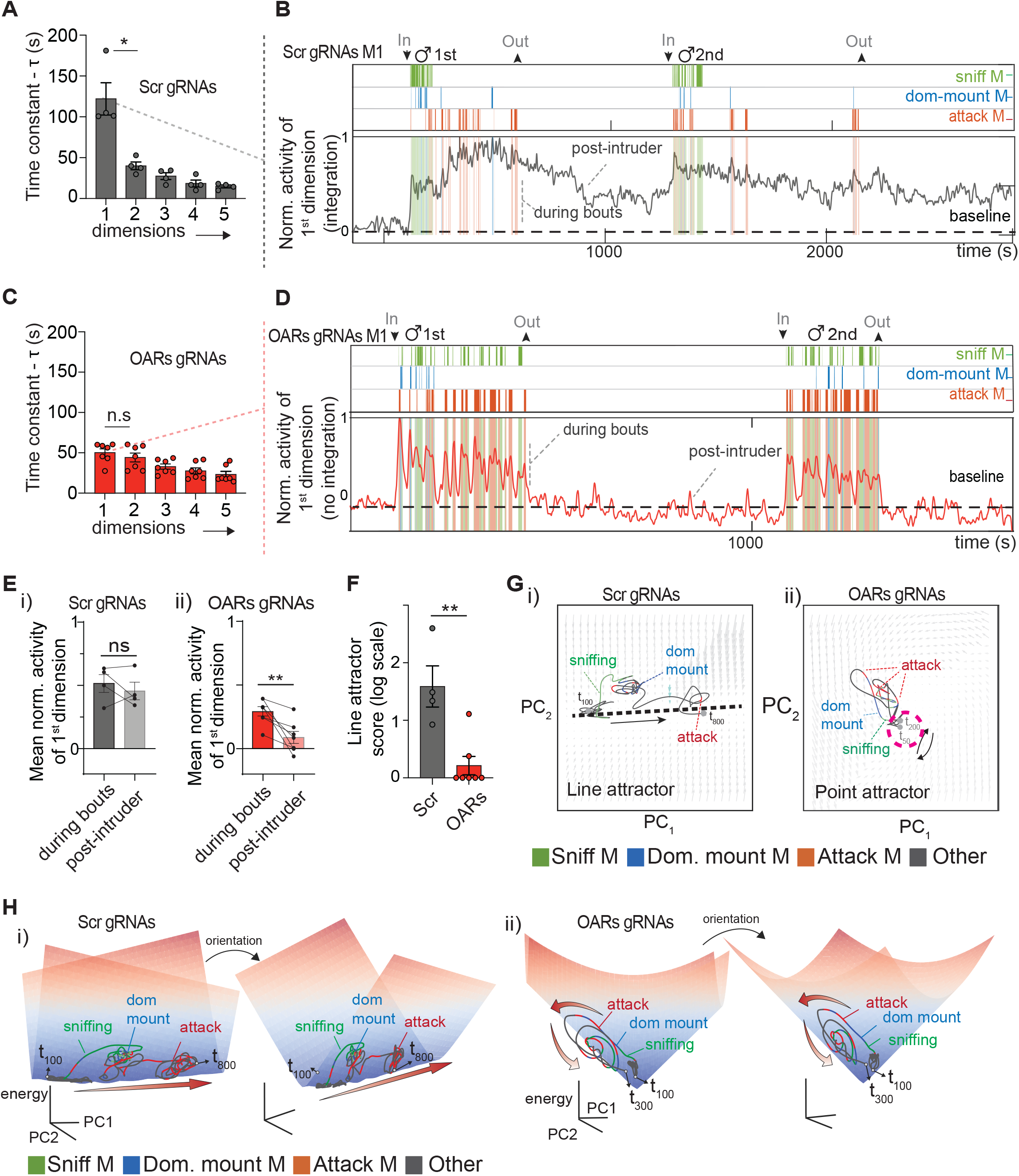
VMHvl^Esr1^ line attractor dynamics require Oxtr/Avpr1a-mediated signaling **A)** Average time constant (τ) of all dimensions, arranged in decreasing order in control mice. **B)** Normalized activity projection onto the time axis of the longest time constant (integration dimension) for VMHvl^Esr1^ units in control mouse M1. **C)** Average time constant of all dimensions, arranged in decreasing order for VMHvl^Esr1^ units in experimental mice. **D)** Normalized activity projection onto the time axis of the 1^st^ dimension (no integration) for VMHvl^Esr1^ units in experimental mouse M1. **E)** Mean normalized VMHvl^Esr1^ activity of the 1^st^ dimension during all behavioral bouts and after removing the intruder male (post-intruder) in control (i) and experimental (ii) mice. **F)** Line attractor score for VMHvl^Esr^ population activity in control and experimental mice. n = 4 control and n =7 in experimental group.**G)** Neural state space with the trajectories projected over time within the inferred flow field rSLDS states for VMHvl^Esr1^ in control (i) and experimental (ii) mouse. **H)** Inferred 3D dynamic landscape in VMHvl control mouse M1(i) and experimental mouse M1(ii). Different views of line (i) and point attractors (ii) are shown. Red arrows depict the neural trajectory associated with attack. 1 minute of sniffing behavior following the introduction of an intruder male is displayed in the behavioral raster plots in (B) and (D). Statistics: Kruskal-Wallis test was performed in (A) and (C). Paired t-test was performed inin (E) and Mann-Whitney test in (F) *p≤0.05 **p≤0.01

To visualize these dynamics in neural state space, we generated 2D flow field graphs spanned by the first two PCs of the fit rSLDS model. The flow fields are comprised of arrows that indicate the rate and direction of change in neural population activity at different points in state space during social interactions (Supplementary Figure S6C). The 2D flow field of control mice (Scr gRNAs) revealed a roughly linear region of low vector flow constituting the line attractor, along which the neural population vector progressed during an inter-male interaction (Figure 6G and Supplemental Figure S6C, dashed black lines). In a 3D dynamic landscape, where the length of the flow-field vectors at each position in neural state space is converted into the height of the landscape (and represented as a heat scale), in control animals neural activity progressed slowly along a trough-like structure (the line attractor) as aggression escalated (Supplementary Figure S6Hi). In contrast, in experimental mice (OARs gRNAs) this line attractor was absent and was replaced by a point attractor (appearing as a white circle in 2D and a cone in 3D), from which the population activity vector made transient excursions during bouts of sniffing or attack (Figure 6G, H and Supplemental Figure S6Cii). This point attractor is a “trivial” fixed point, corresponding to the resting state of a system in which activity decays to baseline in the absence of inputs. These data indicate that Oxtr/Avpr1a-mediated signaling is required for the emergence of line attractor dynamics in VMHvl during aggression.

To investigate in more detail how co-disruption of *Oxtr/Avpr1a* perturbs VMHvl^Esr1^ attractor dynamics, we projected the weighted average of neuronal activity in the 1^st^ rSLDS dimension (which generates the line attractor) onto the time axis and overlayed behavior annotations. As reported previously^20^, in control mice this activity ramped up during the progression from male-directed sniffing to attack, eventually reaching a plateau where it decayed slowly between attack bouts towards a single intruder and remained elevated between sequential trials with different intruders (Figure 6B, Ei and Supplementary Figure S6A, “post intruder”). In contrast, in experimental mice activity in the 1st dimension decayed rapidly in between attack bouts, displaying a “sawtooth” profile, and was significantly lower during the post-intruder (i.e., inter-trial) interval (Figure 6D, Eii and Supplementary Figure S6B). The observation that 1^st^ dimension activity in experimental mice is transiently elevated during aggressive episodes, but does not remain stable across attack bouts and trials, indicates that these neurons are still active during attack, but do not integrate recent experiences in the same way as normal mice.

### Oxtr/Avp1ra-mediating signaling controls VMHvl ^Esr1^ persistent neural activity

We next asked whether changes in the dynamics of VMHvl^Esr1^ neurons caused by co-disruption of *Oxtr/Avpr1a* could account for the elimination of the line attractor in experimental mice. We focused initially on cells that were strongly weighted by the 1^st^ rSLDS dimension (stem plots in Figure 7Aiii). In raster plots from control mice, these units exhibited activity that persisted across inter-attack bout intervals and decayed slowly after removing the intruder male, visible as a “smearing” of rasters over time (Figure 7Ai, Scr gRNAs). In contrast, analogous units from experimental mice exhibited activity time-locked to attack bouts, visible as a vertical stripe-like pattern (Figure 7Ai, OAR gRNAs). To quantify these dynamics, we computed the average autocorrelation half-width (ACHW)^59^, an approximate measure of the decay constant^85,86^, for each 1^st^ dimension-weighted unit. The mean ACHWs during the first minute of interaction were significantly shorter in experimental (9.86 ± 1 s) than in control mice (28.6 ± 1.23 s), by ∼20 seconds (Figure 7Aii, inset).

**Figure 7.**
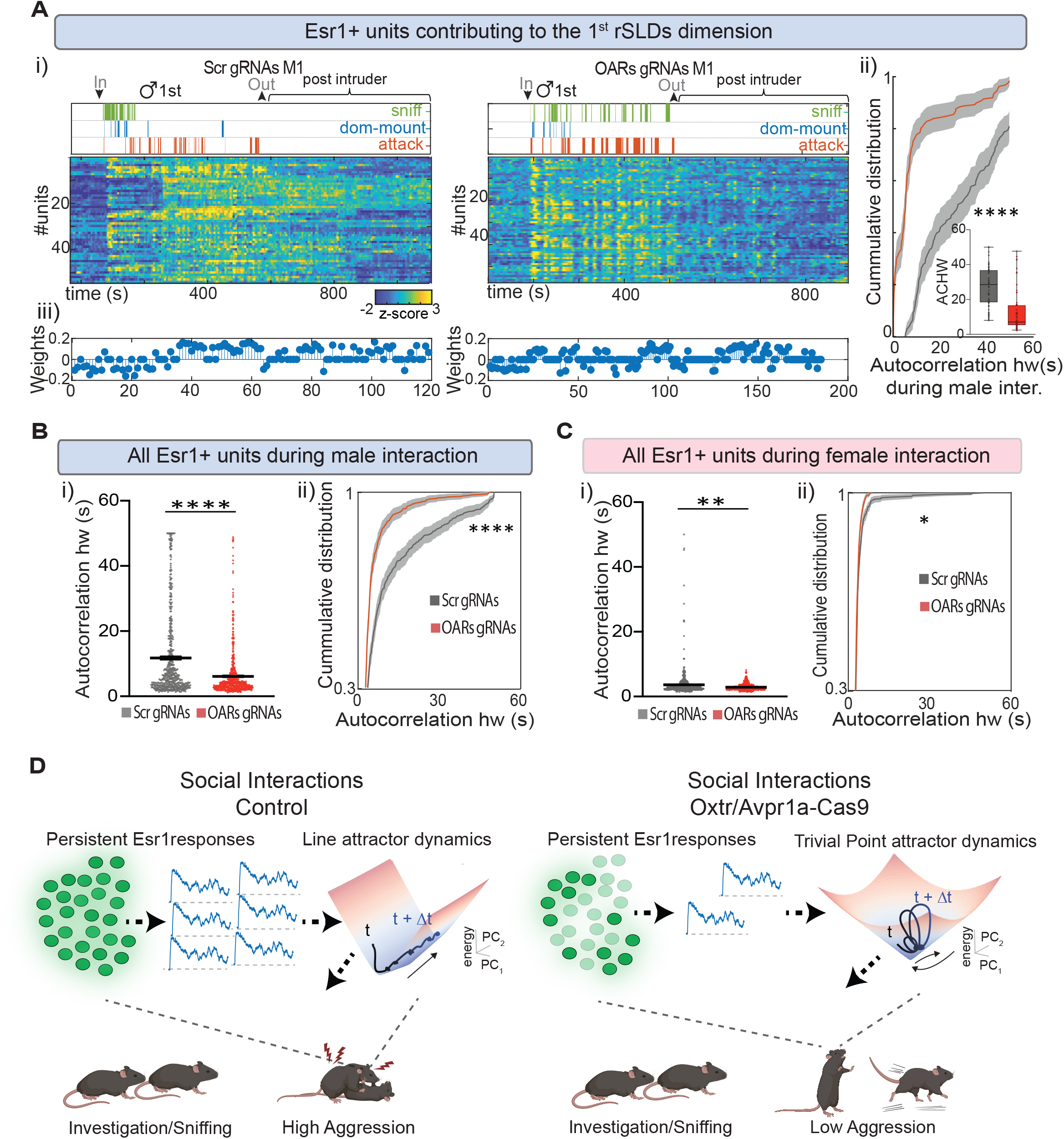
Oxtr/Avp1ra-mediating signaling controls VMHvl ^Esr1^ persistent neural activity. **A)** Behavioral raster plot and the corresponding neural activity (z-score) of individual VMHvl^Esr1^ units that contribute to the 1^st^ dimension in control (left) and experimental (right) mouse (i). Cumulative distribution and inset box plot of VMHvl^Esr1^ persistence, measured by autocorrelation half width (ACHW) (ii) of units identified in the stem plots (iii) from control (n=4) and experimental (n=7) mice. The whiskers in the inset box plot represent the minimum and maximum values. **B)** Average single unit (i) and cumulative distribution (ii) of neuronal persistence measured by ACHW of all VMHvl^Esr1^ recorded units in control and experimental mice during male-male, and C) during male-female interactions. n=5 control, n=7 experimental animals. **D)** A graphic illustration depicting our working model. Oxtr and Avpr1a-mediated signaling control aggression escalation by regulating VMHvl^Esr1^ persistent activity and line attractor dynamics (left) during male-male interactions. However, co-perturbation of *Oxtr* and *Avpr1a* results in a severe reduction of the slow neural dynamics of VMHvl^Esr1^ neurons and the emergence of a trivial point attractor. Under such conditions, male animals fail to display a high level of offensive aggression against male conspecifics (right). Statistics: Values plotted as mean ± S.E.M. Nested Mann-Whitney test was performed for (Bi, Ci, Aii inset). Kolmogorov test was performed (Aii, Bii, Cii) *p≤0.05 **p≤0.01 ****p≤0.0001

A reduction in the average ACHW was also observed in the VMHvl^Esr1^ population as a whole (Figure 7B), although the group difference between means (50.9%) was smaller than that calculated for 1^st^ dimension-weighted units (65.4%). In contrast, the mean and cumulative distribution of ACHWs were only slightly reduced (∼by 22%) during male-female interactions (Figure 7Ci-ii). Thus, the dynamics of individual VMHvl^Esr1^ units are faster, on average, in experimental than in control mice during male-male social interactions. This reduction in ACHWs is consistent with and may explain, the loss of line attractor dynamics caused by co-disruption of *Oxtr/Avpr1a*.

### OXT and AVP evoke persistent responses in VMHvl^Esr1^ neurons *ex vivo*

The observation that co-disruption of *Oxtr/Avpr1a* shortened the average decay time of Esr1^+^ units raised the question of whether adding these peptides to VMHvl would, conversely lengthen this decay. Because it is not technically feasible to apply drugs or peptides directly at the site of miniscope imaging, we utilized the *ex vivo* VMH brain slice preparation (Figure 1F). We transiently perfused the slices with a cocktail of OXT & AVP, imaged activity at single cell resolution, and fit a dynamical systems model to the data using rSLDs (Supplementary Figure S7G). Although this fit model was based on only two dimensions (x1, x2), it captured 85% of the observed variance in neural activity. The time constant of dimension x1 (∼90s) was ∼8-9 -fold greater than that of x2 (∼15s; Supplemental Figure S7H), similar as what we observed *in vivo* (Figure 6A, C). This suggested the existence of two populations of VMHvl^Esr1^ neurons that respond to the neuropeptides with distinct neural dynamics. Indeed, stem plots revealed that the neurons highly weighted by dimension x1 were distinct from those weighted by x2 (Supplementary Figure S7I). Plotting the time-varying average activity of the neurons contributing to these two dimensions revealed that x1 neurons exhibited slowly decaying responses to OXT+AVP application, while x2 neurons exhibited more transient responses (Supplementary Figure S7J-N). These data indicate that acute application of OXT and AVP can evoke persistent responses in a subset of VMHvl^Esr1^ neurons *ex vivo*.

## DISCUSSION

CRISPR/Cas9 gene editing technology^48,87^ has revolutionized virtually every area of biology, but has thus far had relatively little impact on Systems Neuroscience. Using a novel approach that integrates such gene editing^48,87^ with population imaging^49^ and dynamical systems analysis^63^, we show here that OXT and/or AVP receptors are required for persistent neural activity, line attractor dynamics and aggression in VMHvl^Esr1^ neurons that control social behaviors ^54,55,56,58^. More generally, these results demonstrate an important role for peptidergic signaling in population-level slow neural dynamics and attractor formation, and suggest that neuropeptides may control certain behaviors, at least in part, through an influence on such emergent properties. The CRISPRoscopy approach developed here may be generally useful for unifying molecular and circuit-level approaches with “manifold”-level approaches^13,14^ to understand the neural control of behavior, emotion and cognition^22^.

### Methodological considerations

We implemented several technical innovations to combine CRISPR/Cas9-mediated gene editing with calcium imaging in a cell type-specific, anatomically restricted manner. We generated a series of viral vectors, which are distinct from earlier constitutive^88^ or conditional^89^ constructs, to integrate Cre-dependent gene editing with calcium imaging. Our conditional expression system avoids reliance on Cre-dependent chromosomally integrated Cas9 transgenes^90^, eliminating time-consuming and cumbersome genetic crosses, and better aligns GCaMP and Cas9 expression in adult mice, bypassing transient developmental expression of Cre drivers. We also harnessed the multiplex gene editing capabilities afforded by CRISPR/Cas9 to target up to four different genetic loci using a single viral construct. Finally, our system permits the flexible combination of different genetic perturbations with different reporters of neuronal activity or signaling.

### The effects of Oxtr/Avpr1a co-disruption on VMHvl^Esr1^ neuronal activity and behavioral tuning

In principle, the observed reduction in aggressive behavior could be a consequence of a reduction in the activity or attack-specific tuning of VMHvl^Esr1^ neurons. We analyzed the level of activity of these neurons in mice with co-disruption of *Oxtr* and *Avpr1a* using fiber photometry (“CRISPRometry”) and miniscope imaging (“CRISPRoscopy”). A significant reduction in bulk VMHvl^Esr1^ neural activity during attack and sniff episodes was observed in mice with bilateral co-disruption of *Oxtr/Avpr1a.* However, because this manipulation also reduced aggression, those results did not distinguish whether the decreased bulk calcium signal is a cause or consequence of this behavioral phenotype. Unilateral CRISPRoscopy (which does not cause any reduction in aggressive behavior) revealed a small but statistically significant decrease in average unit activity during sniffing or attacking males. This subtle decrease seems unlikely to account for the decreased aggressiveness seen in mice with bilateral gene disruptions, although it formally cannot be excluded.

Previous studies from our laboratory have reported that the percentage of VMHvl^Esr1^ neurons specifically tuned to attack behavior is relatively small, from 2.5-10% ^57–59^. In mice with co-disruption of *Oxtr* and *Avpr1a*, the percentage of attack- and sniff-selective cells was reduced. This result was confirmed by choice-probability analysis^59^ which also revealed an increase in the proportion of cells displaying “mixed selectivity” (i.e., tuned to both sniffing and attack). Whether a relatively large reduction in the small fraction of attack-tuned neurons in VMHvl can account for the behavioral phenotype is not yet clear, especially if some of those cells are converted to mixed-selectivity neurons, rather than simply inactivated. Previous studies have shown that attack behavior can be accurately decoded from VMHvl^Esr1^ population activity despite the dearth of attack-tuned cells^59^, and we show here that the performance of such decoders was not affected by co-disruption of *Oxtr/Avpr1a*.

### The impact of Oxtr/Avpr1a co-disruption on neural dynamics

More striking physiological phenotypes were observed in experimental mice when we examined the dynamics of VMHvl^Esr1^ neuronal activity. In control mice many of these neurons exhibited persistent activity, with autocorrelation half-widths (ACHWs; an approximation of the neuronal time constant) ranging over tens of seconds. Co-disruption of *Oxtr/Avpr1a* caused a strong and highly statistically significant reduction in the average ACHW of VMHvl^Esr1^ neurons. These data reveal a requirement for normal OXTR/AVPR1a-mediated signaling in persistent neural activity at the population level. Our between-subject comparisons do not allow us to distinguish whether co-disruption of *Oxtr/Avpr1a* reduced the population average time constant by changing the dynamics of individual cells, or simply by reducing the proportion of active cells with fixed, slow dynamics. However, bath application of OXT+AVP induced persistent calcium responses in a subset of VMHvl^Esr1^ neurons *ex vivo* (Supplemental Figure S7). Interestingly, persistent activity in entorhinal cortical neurons has previously been shown to require muscarinic cholinergic signaling *ex vivo*^82^, and is thought to underlie some forms of working memory^83,91^.

Consistent with these changes in single unit dynamics, fitting a linear dynamical systems model^63^ to the CRISPRoscopy data revealed a large decrease in the time constant of the 1^st^ rSLDS dimension. Accordingly, the average time-varying activity of these neurons did not show persistence during aggression, but rather exhibited a “saw-tooth”-like pattern time-locked to individual attack bouts. In addition, persistent activity between aggression trials^20^ was severely reduced. Most strikingly, in 2D and 3D projections of the rSLDS manifold the line attractor^20^ was absent and replaced by a point attractor centered on the resting state (Figure 6F, G).

A parsimonious hypothesis is that the reduction in single-unit persistent activity caused by co-disruption of *Oxtr/Avpr1a* is responsible for the loss of line attractor dynamics. These changes in dynamics may in turn be responsible for the reduction in aggressive behavior that is caused by the same perturbation. However, a contribution from the small but significant reductions observed in the number, activity and tuning specificity of attack-selective neurons cannot be excluded.

### A line attractor dependent on neuropeptide signaling

The data presented here provide to our knowledge the first evidence in any system of a requisite role for neuropeptide signaling in line attractor dynamics. In contrast, most theoretical studies of attractor dynamics to date have assumed that they reflect recurrent fast synaptic connectivity^17,92^, although that has been experimentally demonstrated so far only in the *Drosophila* ring attractor system^23,28^. The finding that neuropeptide signaling is required *in vivo* for persistent activity and line attractor dynamics in VMHvl is consistent with recent slice recording studies reporting sparse local fast glutamatergic connectivity and predominantly neuropeptidergic transmission in this nucleus^93^. Nevertheless, the two mechanisms are not mutually exclusive. Indeed, in an earlier study we generated recurrent network models of persistent activity displayed by VMHdm^SF1^ neurons that mediate defensive behaviors^94–96^, and found that only models in which local transmission occurred on a time scale consistent with slow neuromodulatory signaling were consistent with our observational data^11^.

The finding that a line attractor which represents an internal motive state of aggressiveness^20^ is dependent on neuropeptidergic signaling is important for several reasons. First, neuropeptide gene expression is strongly modulated by physiological influences including hormones^97^ and neural activity^98^. Through this mechanism, such influences may mediate physiological state-dependent changes in attractor dynamics^19^. Second, neuropeptide receptor expression is more restricted than the expression of receptors for biogenic amines^99^, another important class of neuromodulators. This specificity could allow the regulation of different (and potentially competing) attractors within the same network by distinct neuropeptides, depending on differential receptor expression; VMH neurons collectively express dozens of neuropeptides and close to 100 GPCRs^69,100^. Finally, neuropeptides such as CRH^101^ and NkB^30,102^ are known to mediate responses to psychological stressors, which are known etiological factors in some neuropsychiatric disorders. It has been proposed that these disorders may reflect changes in attractor dynamics in certain brain regions^103^. Our data suggest that such changes could potentially result from stress-induced alterations in neuropeptide signaling^104,105^ that is necessary for attractor formation.

The results described here constitute a “first-in-class” experiment that demonstrates, on the one hand, how the power of multiplex gene editing technology^48,87^ can combined with single cell imaging to link emergent properties discovered by dynamical systems analysis to underlying molecular control mechanisms^17^. On the other hand, they open a new window onto systems-level mechanisms that may mediate the behavioral functions of some neuropeptides^39,41^. In this way, this work may help to enable mechanistic explanations in neuroscience that unify different levels of abstraction and of biological organization^22^.

### Limitations

A technical limitation of our approach is that two different viruses are used to deliver gRNAs-GCaMP and Cas9. Consequently, not all GCaMP^+^ cells captured in our imaging analysis are necessarily Cas9^+^ (and therefore mutant for *Oxtr/Avpr1*). Our data indicate co-infection rates of ∼65-80% depending on the viruses used. Developing single-vector approaches^106^ would ensure that calcium imaging is restricted to cells that co-express both Cas9 and gRNAs. Another limitation is that we lack an accurate estimation of the fraction of cells exhibiting homozygous INDELs in the *Oxtr* and *Avpr1a* loci *in vivo*. Consequently, our results may underestimate the magnitude and penetrance of the effects of a complete null mutation of *Oxtr/Avpr1a* in all VMHvl^Esr1^ neurons. Another limitation is that our statistics are based on between-subject comparisons of experimental vs. control mice, necessitating large sample sizes for behavioral experiments. Statistical power could be increased by using an inducible CRISPR/Cas9 system permitting within-subject comparisons, which would also clarify the interpretation of the underlying mechanism. Finally, our results do not distinguish the respective roles of OXTR and AVPR1a, nor do they establish the cellular source and identity of the endogenous peptide(s) release that activates these receptors *in vivo* during aggression. Further studies will be required to elucidate these biologically important details.

## ACKNOWLEDGEMENTS

We thank H. Inagaki, A. Kennedy, B. Weissbourd, M. Schnitzer and S. Linderman for critical feedback on this manuscript, C. Chiu for laboratory management, G. Mancuso and L. Chavarria for administrative assistance, and members of the Anderson lab for helpful comments on this project. G.M. has been supported by post-doctoral fellowships from the Tianqio and Chrissy Chen Institute for Neuroscience, the Helen Hay Whitney Foundation and the Della Martin Foundation. A. N. is supported by a National Science Scholarship from the Agency of Science, Technology and Research, Singapore. This work was supported in part by NIH grants NS123916, MH1223612, and MH070053. D.J.A. is an Investigator of the Howard Hughes Medical Institute. The content of this paper is solely the responsibility of the authors and does not necessarily represent the official views of the National Institutes of Health. We support inclusive, diverse, and equitable conduct of research. We acknowledge that this research was conducted at Caltech, which is located on the unceded land of the Indigenous Tongva people.

## Author contributions

G.M and D.J.A conceived the project. G.M and D.J.A. wrote and prepared the manuscript, with input and help from A.N. G.M performed all the experiments, with the occasional help from B.Y for the miniscope imaging and D.K. for the two-photon slice imaging. G.M and A.N performed quantitative measurements of behavior and microendoscope imaging data. A.N performed the dynamical system modeling and decoder analyses.

## SUPPLEMENTAL FIGURE LEGENDS

**Figure S1. Expression and *ex-vivo* functional characterization of *Oxtr* and *Avpr1a* in the VMH, related to Figure 1**

**A)** A diagram portraying the diverse methodologies employed to investigate the role of neuromodulators in the mammalian brain. Among the most recent predominant strategies are the systemic administration of agonists/antagonists targeting G-protein-coupled receptors (GPCRs)^107^ or the use of knock-out animals combined with *in vivo* single recording unit approaches^108^. However, there is a requirement for novel regionally restricted gene-editing strategies, with cell type specificity^109^, compatible with bulk^88,110,111^ or single-cell calcium recording methods^90,112^ from many cells in freely moving animals (Mountoufaris et al). **B)** Violin plots illustrating the expression of *esr1, avpr1a,* and *oxtr* mRNAs outside the VMH and in non-neuronal clusters; “max CPM”, maximum counts per million reads. **C)** t-SNE plots illustrating the distribution of *esr1* (i) or *avpr1a* and *oxtr* mRNAs (ii; from Figure 1C) in single cells from the VMH. “FPKM”, fragments per kilobase of exon per million mapped fragments. **D)** Violin plots illustrating the expression of *Esr1, Oxt,* and *Avp* mRNAs in VMHvl; “max CPM”, maximum counts per million reads. **E)** Venn diagram of *esr1, avpr1a,* and *oxtr* mRNA expressing neurons in the ventral part of the VMH and the surrounding areas. **F)** Quantification of Esr1^+^ responses to 400nM of Avp (n=62 cells from 3 mice) or OXT (n=79 cells from 3 mice). n=70 cells not activated in brain slices from 3 mice. The shadow area depicts the duration of the bath application of peptides. **G**) An example field of view of spontaneous active (left) and induced Esr1^+^ GCaMP7f calcium responses to 400nM OXT & AVP. Statistics: Nested Mann-Whitney test was performed, and values were plotted as mean ± S.E.M. in (D). ****p≤0.0001

**Figure S2. Experimental validation and behavioral characterization of CRISPR/Cas9 based *Oxtr* and *Avpr1a* perturbations, related to Figure 2**

**A)** T7E1-endonuclease treated *Oxtr* (left) and *Avpr1a* (right) PCR products from N2a-Cas9 expressing cells, transfected with gRNAs against *Avpr1a or Oxtr*. As a negative control, scrambled gRNAs (Scr) were used. Yellow stars indicate the cleavage products, indicating successfully induced Cas9-mediated insertions or deletions (INDELs) in the coding sequence complementary to the gRNAs. **B)** Quantification of the percentage of animals attacking (i), the average duration of each attack bout (ii), the latency of the 1^st^ attack bout (iii), and the interval between attack bouts (iv) against a male intruder. n=11 mice per group. **C)** Quantification of the number and time-varying probability of attack bouts against a male intruder (i-ii). n=11 mice per group. Quantification of the average velocity during attack in control (n=9 mice) and experimental mice (n=8 mice) (iii). **D)** Quantification of the percentage of animals mounting (i), the average duration of each mounting bout (ii), the latency of the 1^st^ mounting bout (iii), and the interval between mounting bouts (iv) in control and experimental mice during male-female interactions. n=11 mice per group. Statistics: nested Mann-Whitney test was performed, and values were plotted as mean ± S.E.M. *p≤0.05. ****p≤0.0001.

**Figure S3. Effects of *Oxtr* and *Avpr1a* co-perturbation on male-, female-directed behaviors and bulk calcium activity, related to Figure 3**

**A)** Immunostaining against the GCamp8s (green) and Cas9 (red) proteins in coronal hypothalamic sections from ESR1-2A-CRE animals co-injected with a Cre-dependent Cas9 AAV and Cre-dependent gRNA-Gcamp8s AAV. Sections counterstained with Dapi (blue). Arrows depict the expression of Cas9 protein in GCamp8s Esr1 double-positive cells (i). Quantification of GCamp8s positive cells expressing the Cas9 protein (ii). n= 3 mice. **B)** Quantification of the number of Esr1^+^ neurons responding to 400nM of AVP and OXT in brain slices from ESR1-2A-CRE mice co-injected with the Cas9 AAV and Cre-dependent OARs-GCaMP8s AAV (experimental group) or Cre-dependent Scr-GCaMP8s AAV (control group) in the VMHvl (see Figure 3Ai). Data points represent brain slices. n=2 Scr RNA (control) and n=3 OARs (targeted) mice. **C)** Quantification of the percentage of animals attacking (i), the average duration of each attack bout (ii), the latency of the first attack (iii), the interval between attack bouts (iv), the number and the time-varying probability (iv-v) of attack bouts during male-male interactions. n=16 control and n=13 experimental mice. **D)** Quantification of the average duration of each mounting bout (i), the latency of the first mount (ii) and the interval between mounting bouts (iii) during male-female interactions. n=15 mice each group. **E)** Z-scored BTA of VMHvl^Esr1^ activity (i) and the area under the curve (ii) during female-oriented sniffing (1 min). n=4 control and n=5 experimental mice. **F)** Z-scored BTA of VMHvl^Esr1^ activity (i) and the area under the curve (ii) during mounting. n=3 control and n=4 experimental mice. Statistics: Mann-Whitney test was performed, and values were plotted as mean ± S.E.M. **p≤0.01. ****p≤0.0001.

**Figure S4. Effect of Oxtr/Avpr1a co-editing on intruder sex-specific representations, single unit neural activity and tuning, related to Figure 4**

**A)** VMHvl^Esr1^ ensemble representations of intruder sex, for control (i) and experimental (ii) mice, projected onto the first two axes of a PLS regression against intruder sex. Traces are colored by intruder sex identity. The percentage of variance explained by the first two PLS components is noted for each male resident. **B**) Quantification of the PLS1 variance explained (which accounts for intruder sex) in control and experimental mice. **C)** Cumulative distribution of z-scored activity of all VMHvl^Esr1^ units during 1 min interaction with male (i) or female (ii) intruders in control and experimental mice. **D)** Average single unit activity (σ) of male responses (ý 2σ relative to the pre-intruder baseline) between control and experimental mice. **E)** Percentage of male- or female selective or co-active units (ý 2σ relative to the pre-intruder baseline), per imaged control (i) or experimental (ii) mouse. n=5 control, n=7 experimental animals. Statistics: Values in (C) are plotted as mean ± S.E.M. Nested Mann-Whitney test was performed in (B) and (D), while nested Kolmogorov–Smirnov test was used in (C) **p≤0.01 ***p≤0.001

**Figure S5. Effect of Oxtr/Avpr1a co-editing on male-, female-directed single unit activity and tuning, related to Figure 5**

**A)** Scatter plot of the average single VMHvl^Esr1^ unit activity (z-score) during male (i) and female (ii) directed behaviors in control and experimental mice. **B)** Cumulative distribution of VMHvl^Esr1^ activity (z-score) during male-directed (sniffing and attack; i & ii) and female-directed behaviors (sniffing and mounting; iii-iv) in control and experimental mice. **C)** Average z-scored VMHvl^Esr1^ activity normalized to the pre-behavior bout period during male-directed sniffing (i), attack (ii), female-directed sniffing (iii) and mounting (iv). **D)** Choice probabilities histograms of male-directed behaviors (attack vs. sniffing) in control and experimental mice. Mixed-tuned units (blue) and units tuned for other behaviors (yellow) are highlighted. **E)** Accuracy of frame-wise decoders predicting attack vs sniffing (i) and attack vs no attack (ii) trained on VMHvl^Esr1^neural activity in control or experimental animals. Decoders were trained and tested on held out data from each group separately. Statistics: Values plotted as mean ± S.E.M. in (B). Nested Kolmogorov–Smirnov test was used in (B), whereas nested Mann-Whitney test was performed in (C) and (E). *p≤0.05 **p≤0.01 ****p≤0.0001

**Figure S6. VMHvl^Esr1^ line attractor dynamics require Oxtr/Avpr1a-mediated signaling, related on Figure 6**

**A)** Normalized activity projection onto the time axis of the longest time constant (integration dimension) for VMHvl^Esr1^ units in control and **B)** in experimental mice. **C)** Schematic illustrating inferred dynamics shown as flow fields, with the line attractor illustrated by a black dashed line in control (i) and the point attractor with a purple dashed circle in experimental mouse (ii). The different rSLDS states (S1-S3) are depicted with different colors. **D)** Quantification of the average time constant (τ) of the 1^st^ dimension in control and experimental mice. n=4 control and n=7 mutant. 1 minute of sniffing behavior following the introduction of an intruder male is depicted in the behavioral raster plots in (A) and (B). Statistics: Mann-Whitney test was performed in (D).

**Figure S7.**

**A)** rSLDs modeling of Esr1^+^ cells calcium responses to 400nM OXT & AVP. Acute brain slices from male ESR1-2A-CRE animals that express Cre-dependent GCaMP7f were used. **B)** Average time constant of the identified two dimensions (X_1_, X_2_), arranged in decreasing order for acute brain slices. **C)** Absolute rSLDS weight of neurons contributing to X_1_ (top) and X_2_ (bottom) dimensions sorted by choice probability values for activity during the OXT & AVP perfusion window. **D)** Heat maps of OXT & AVP induced slow persistent X_1_ (left) and **E)** transient X_2_ responses (right) VMHvl^Esr1^ neurons. **F)** Projection of population activity onto the time axis of the slow persistent X_1_ and **G)** the transient X_2_ dimensions. **H)** Overlay projection of population activity onto the time axis of the slow persistent X_1_ and the transient X_2_ dimensions.

## STAR METHODS

### EXPERIMENTAL MODEL AND SUBJECT DETAILS

All procedures were performed in accordance with NIH guidelines and approved by the Institutional Animal Care and Use Committee (IACUC) at the California Institute of Technology (Caltech). We used *Esr1^Cre/+^* ^55^ and *Oxtr ^Cre/+^* ^113^ transgenic mice. Animals were housed and maintained on a reverse 12 h light-dark cycle with food and water ad libitum. We used wild-type (WT) C57BL/6N male mice (experimental), C57BL/6N female mice or BALB/c females (for sexual experience), and BALB/c male mice (intruders) were obtained from Charles River (Burlington, MA). Behavior was tested during the dark cycle.

#### Viruses

The following <u>AAVs</u> were used in this study, with injection titers as indicated. Viruses with a high original titer were diluted with clean PBS on the day of use. AAV1-Syn-Flex-GCaMP7f (2.1 x e^13^) was purchased from Addgene. AAVDJ/8-EFS-NC-SpCas9-HA -NLS-Poly(A) (pBK694) (2.00x e^13^) and AAV9-EFS-NC-SpCas9-HA -NLS-Poly(A) (2.17x e^13^) purchased from Duke viral vector core. The AAV9-EFS-NC-DIO-SpCas9-myc-NLS-Poly(A) (2.31x+e^13^), AAV9-hPGK-DIO-SpCas9-myc-NLS-Poly(A) (2.31x+e^13^), AAV1-4xgRNAs_scramble_-hSyn flex-GCaMP8s-wpre (2.75x e^12^) and AAV1-4xgRNAs _Oxtr/1Avp1ra_ -hSyn flex-GCaMP8s-wpre 2.44x e^12^) were packaged at the HHMI Janelia Research Campus virus. The 4xgRNAs_scramble_-ubc DsRed (TU/ML 1.5X e^8^) and 4xgRNAs_Oxtr/Avp1ra_ ubc DsRed (OXTR/AVP1Ra) lentiviruses (TU/ML 3.50 X e^8^) were also packaged at the HHMI Janelia Research Campus virus facility.

## METHOD DETAILS

### Generation of multiplex CRISPR/*Streptococcus pyogenes* Cas9 vectors

The multiplex gRNAs construct was based on the backbone of pLV GG hUbC-dsRED (addgene #84034) after being modified to remove superfluous sequences (e.g., LoxP). Individual gRNAs were cloned as described in *Kabadi et al., 2014*^73^. Cas9 pAAV-EFS-NC-SpCas9-NLS-Poly(A) (Duke viral vector core) vectors were used for constitutive Cas9 expression. For Cre-dependent CRISPR/Cas9 DIO/FLEX, sequences were cloned in pAAV-EFS-NC-Cas9-NLS-Poly(A) vectors, respectively. The Golden Gate cassette to express four gRNAs was cloned into the pGP-AAV-syn-FLEX-jGCaMP8s-WPRE (addgene 162377)^114^ vector. Four different gRNAs are constitutively being expressed under four different polymerase llI promoters, and a Cre-dependent expression of the calcium indicator jGCaMP8s under the express human synapsing promoter.

### Generation of Neuro2A constitutively expressed CAS9 cells and screening of gRNAs

Neuro2a cells (ATCC CCL-131^™)^ were transfected with EFS-Cas9-Blast plasmid (addgene # 52962) and selected for four days with blasticidin antibiotic. Individual gRNAs were cloned in ph7SK-gRNA (addgene # 53189) or phH1-gRNA (addgene # 53186) or pmU6-gRNA (addgene # 53187) or phU6-gRNA (addgene # 53188) plasmids, 0.5-1 ug transfected in Neuro2a-Cas9 cells and after three days genomic DNA was isolated. We used T7 endonuclease assay for testing the efficiency of each gRNAs to generate edits, similar to what has been described in https://www.neb.com/protocols/2014/08/11/determining-genome-targeting-efficiency-using-t7-endonuclease-i. In addition, Sanger traces were generated with target-specific PCR and analyzed with the Tracking of INDELs by Decomposition (TIDE) web tool http://tide.nki.nl (data not shown).

### Screening for aggressor male and resident intruder assay

All experimental male mice (‘‘residents’’) were individually housed for two weeks and received sexual experience (for at least one week). Previously it has been reported that ∼20-25% of inbreeding C57BL/6N male animals fail to display territorial aggression against conspecific male intruders during the RI assay ^115^. We pre-screened males for baseline aggression using resident-intruder testing sessions to identify and exclude no aggressors from our analysis. Animals that attacked two constitutively presented intruders were termed aggressors and added to the pool of animals for CRISRP/Cas9-based gene editing surgeries. On the experimental day, the preselected male residents were transported in their home cage to a novel behavioral testing room (under infrared light), where they acclimated for 5-10 min. An unfamiliar group housed BALB/c mouse (‘‘intruder’’) was then placed in the resident’s home cage, and residents were allowed to interact with it for period of time.

### Acute brain slices preparation

Briefly, male adult mice were anesthetized with isoflurane and transcardially perfused with cold NMDG-ACSF (adjusted to pH 7.3–7.4) containing CaCl2 (0.5 mM), glucose (25 mM), HCl (92 mM), HEPES (20 mM), KCl (2.5 mM), kynurenic acid (1 mM), MgSO4 (10 mM), NaHCO3 (30 mM), NaH2PO4 (1.2 mM), NMDG (92 mM), sodium L-ascorbate (5 mM), sodium pyruvate (3 mM), thiourea (2 mM), bubbled with carbogen gas (95% O_2_ and 5% CO_2_). The brain was sectioned at 250 μm using a vibratome (VT1000S, Leica Microsystems) on ice and was incubated in 34oc for 12 min, in NMDG – ACSF. Then transfer the sections to room temperature in aCSF/HEPES-GSH solution (adjusted to pH 7.3–7.4,) containing CaCl2(2 mM), glucose (25 mM), kCl (2.5 mM), HEPES (20 mM), NaCl (92 mM), MgSO4 (2 mM), NaHCO3 (30 mM), NaH2PO4 (1.2 mM), sodium L-ascorbate (5 mM), sodium pyruvate (3 mM), thiourea (2 mM), and Glutathione Monoethyl Ester (0.5-1mM)-before proceeding with Ca^2+^ imaging.

### Peptide perfusion and two-photon calcium imaging experiments

Solutions of 100nM or 400nM of Oxt, Avp, and Oxt + Avp peptides were prepared in aCSF and perfused with a rate of 1-2ml/min through a microfluidics chamber containing the brain slices. Calcium imaging was performed using a custom-modified Ultima two-photon laser scanning microscope (Bruker). The primary beam path was equipped with galvanometers driving a Chameleon Ultra II Ti:Sapphire laser (Coherent) and used for <u>GCaMP</u> imaging (920 nm). GCaMP emission was detected with photomultiplier-tube (Hamamatsu). Images were acquired with an Olympus20X XLUMPLFLN Objective, 1.00 NA, 2.0 mm WD. All image acquisition was performed using PrairieView Software (Version 5.3) with a framerate of ∼1.2Hz.

### Behavior recording

All behavioral experiments were performed in conventional mouse housing cages (home cage or new cage) under red lighting, using the previously described behavior recording setup ^116^. The behavior video’s top and front views were acquired at 30 Hz using the video recording software, StreamPix7 (Norpix).

### Behavior annotations

Behavior videos were processed using an automated behavior classification system to generate frame-by-frame annotations of attack, mounting and sniffing behavior ^117^. The output of the classifier and behavior videos were loaded into a MATLAB based MATLAB-based behavior annotation interface and then manually corrected by trained individuals to produce a final set of annotations^117^. A ’baseline’ period of 5-minutes was recorded at the start of every recording session during which the animal was alone in its home cage. Six behaviors were annotated during the resident intruder assays: sniff (face, body, genital-directed sniffing), towards male or female intruders, and attack, mount with male or female intruders. For quantifying the interval (s) between behavioral bouts in Figures 2, 3, S2 and S3 animals that didn’t show any mount or attack behavior were excluded. 15 min for male-male interaction was scored in Figure 2 and S2. In Figure 3 and S3 ∼20 min of RI was scored for male -male interaction. 10 min of male-female interaction was scored during the RI assays in Figure 2, 3, S2 and S3.

In addition to the classification of behaviors, automated pose estimation was performed on behavior videos to obtain key points of interacting mice^117^. The velocity of the resident mouse was calculated as the change in positions of centroids of the head and hips, computed across two consecutive frames as previously performed^58,117^. The distribution of this feature was computed for both experimental and control animals to obtain the data shown in Supplementary Figure S2C(iii).

### Stereotaxic surgery

Surgeries were performed on socially and sexually experienced adult male *Esr1^Cre/+^* mice and *Oxtr ^cre/+^* mice 8–12 weeks old. Virus injection and implantation were performed as described previously ^58,59^. Briefly, animals were anesthetized with isoflurane (5% for induction and 1.5% for maintenance) and placed on a stereotaxic frame (David Kopf Instruments). The virus was injected into the target area using a pulled-glass capillary (World Precision Instruments) and a pressure injector (Micro4 controller, World Precision Instruments) at a 20 nl /min flow rate. The glass capillary was left in place for 5 -10 minutes following injection before withdrawal. The injection volumes were ∼400-500nl for bilateral injection in mice used for behavioral analysis and CRISRPometry. For micro endoscope recordings, we performed unilateral ∼200nl injections. The Stereotaxic injection coordinates were based on the Paxinos and Franklin atlas (VMHvl, anterior– posterior: −4.68, medial–lateral: ±0.73, dorsal–ventral: −5.73). For single fiber optogenetic and fiber photometry experiments (optogenetics: diameter 200 μm, N.A., 0.22; fiber photometry: diameter 400 μm, N.A., 0.48; Doric lenses) were then placed above the virus injection sites (fiber photometry: 150 μm above) and fixed on the skull with dental cement (Metabond, Parkell). For micro-endoscope experiments, virus injection and lens implantation were performed on the same day Lenses with a baseplate were slowly lowered into the brain and fixed to the skull with dental cement. Mice were habituated with weight-matched dummy micro-endoscopes (Inscopix) for at least one week before behavior testing. Mice were head-fixed on a running wheel 3-4 weeks after lens implantation, and a miniaturized micro-endoscope (nVista, Inscopix) was attached to the baseplate for imaging. Mice were singly housed after surgery and were allowed to recover for at least 4 weeks before behavioral testing.

### Histology

Once the behavioral experiments were finished, virus expression and implant placement were histologically verified on all mice. Mice lacking correct virus expression or implant placement were excluded from the analysis. Mice were transcardially perfused with 1x PBS at room temperature, followed by 4% paraformaldehyde (PFA) (diluted from 16% EM grade PFA). Brains were extracted and post-fixed in 4% PFA 16-24h at 4°C, followed by 24 hours in 30% sucrose/PBS at 4 °C. Brains were embedded in OCT mounting medium, frozen on dry ice and stored at −80°C for subsequent sectioning. Brains were sectioned into 60 μm slices on a <u>cryostat</u> (Leica Biosystems). Sections were washed with 1× PBS and mounted on Superfrost slides, then incubated for 15 minutes at room temperature in DAPI/PBS (0.5 μg/ml) for counterstaining, rewashed and coverslipped. Sections were imaged with an epifluorescent microscope (Olympus VS120) For some epitope staining 30 um sections were cut from either fresh-frozen tissue or post-fixed 2h 4%PFA on ice, immersed in 30% sucrose:1xPBS 4C 2h before embedding in OCT. For Cas9 immunostaining, a cocktail of antibodies against the epitope and the Cas9 protein was used. Animals were stained after 9-12 weeks post-injection.

### Fiberphotometry recordings

The fiber photometry setup was similar to what was previously described ^118^. We used 470 nm LEDs (M470F3, Thorlabs, filtered with 470-10 nm bandpass filters FB470-10, Thorlabs) for fluorophore excitation and 405 nm LEDs for isosbestic excitation (M405FP1, Thorlabs, filtered with 410–10 nm bandpass filters FB410-10, Thorlabs). LEDs were modulated at 208 Hz (470 nm) and 333 Hz (405 nm) and controlled by a real-time processor (RZ5P, Tucker David Technologies) via an LED driver (DC4104, Thorlabs). The emission signal from the 470 nm excitation was normalized to the emission signal from the isosbestic excitation (405 nm), to control for motion artifacts, photobleaching, and levels of GCaMP8s expression. LEDs were coupled to a 425 nm longpass dichroic mirror (Thorlabs, DMLP425R) via fiber optic patch cables (diameter 400 μm, N.A., 0.48; Doric lenses). Emitted light was collected via the patch cable, coupled to a 490 nm long pass dichroic mirror (DMLP490R, Thorlabs), filtered (FF01-542/27-25, Sem-rock), collimated through a focusing lens (F671SMA-405, Thorlabs) and detected by the photodetectors (Model 2151, Newport). Recordings were acquired using Synapse software (Tucker Davis Technologies). On the test day, after at least 5 minutes of acclimation under the recording setup, the male resident was first recorded for 1 minute to establish a baseline. Male or female intruders were introduced into the home cage on separate days. Typically, each session lasted 15-20 min.

### Microendoscope recordings

On the day of imaging, mice were habituated for at least 5-10 minutes after installing the micro endoscope in their home cage before the start of the behavior tests. Imaging data were acquired at 30 Hz with 2× spatial down sampling; light-emitting diode power (0.1–0.5) and gain (1–7×) were adjusted depending on the brightness of GCaMP expression as determined by the image histogram according to the user manual. A transistor–transistor logic (TTL) pulse from the Sync port of the data acquisition box (DAQ, Inscopix) was used for synchronous triggering of StreamPix7 (Norpix) for video recording. Imaging sessions typically lasted 1 h (20–25 min interactions per sex).

### Micro-endoscopic data extraction

Preprocessing and Calcium data extraction was performed similarly to what has been previously described ^57^. Briefly, data were 2x downsampled, motion corrected, and a spatial band-pass filter was applied to remove the out-of-focus background. Next, filtered imaging data were temporally downsampled to 10 Hz. Calcium traces were extracted and deconvolved using the CNMF-E ^119^ with the following parameters: patch_dims = [42, 42], gSig = 3, gSiz = 13, ring_radius = 19, min_corr ∼0.57-0.62, min_pnr = ∼5.5-6, deconvolution: foopsi with the ar1 model<u>44</u>. Every extracted unit’s spatial and temporal components were manually inspected (SNR, PNR, size, motion artifacts, decay kinetics, etc.). Traces of units were either z-scored or normalized in units of σ relative to the baseline fluorescence (during 7sec or more) of the neuron before the first trial of resident-intruder interactions, as previously described^57,58^, Distinct hypothalamic control of same-and opposite-sex mounting behavior in mice. In Supplementary Figure 5 C, the z-scored value during a behavioral bout for each unit was normalized by subtracting the mean of a 2-3 sec baseline before the onset of the bout. The average normalized activity was quantified for a period of 15 sec. A total of 585 units (n=5 mice) from control and 546 units (n=7 mice) from experimental mice were recorded.

### Quantification and statistical analysis

#### Transcriptomic analysis of *Esr1*, *Oxtr*, *Avpr1a* mRNAs

The violin plots, t-SNE and the quantification of *Esr1*, *Oxtr*, *Avpr1a* mRNAs in the different single cell VMH clusters (Figure 1B, S1B and S1D) or in single Esr1+ cells (Figure 1D) was performed with R package Seurat as described previously^69^.

#### Fiberphotometry analysis

All data analyses were performed in Matlab 2020a and Python 3.8.3 as previously described^58^. Briefly, behavioral video files and fiber photometry data were obtained in a time-locked manner. Photometry recordings yielded both a 405-nm (isosbestic, Ca^2+^ independent) signal and a 470-nm (Ca^2+^ dependent)) signal. To align the 405-nm signal to the 470-nm signal, a least squares linear fit is first performed. The motion corrected 470-nm signal is obtained as follows: [*F*_470_(t) – *F*_405_(*t*)] / *F*_405_(*t*). To normalize activity, the baseline value F_0_ and standard deviation SD_0_. were calculated using a 2 second window as follows [*F*_*n*_(*t*) – *F*_0_(*t*)] /*SD*_0_. Overlapping behavioral bouts within this time window were excluded from the analysis. The peak and area under the normalized activity curve (AUC) were calculated within the 10-second window. We confirmed that the latency to achieve the peri-stimulus time histogram (PETH) peak level is shorter than the indicated time window.

### Miniscope neural data analysis

#### Choice probability

Choice probability (CP) analysis was used as before^58^ to measure a cell’s tuning, defined here as how well two conditions could be predictively discriminated from a single cell’s activity ^120^. The CP of a given cell for a pair of behavioral conditions was computed by constructing a histogram of that cell’s Δ*F*(*t*)/*F*_0_ values under each of the two conditions. These two histograms were plotted against each other to generate a ROC (receiver-operating characteristic) curve. The integral of the area under this ROC curve generated the CP value for each cell with respect to each of the two behavioral conditions. This CP value is bounded from 0 to 1, where a CP of 0.5 indicates that the neuron’s activity cannot distinguish between the two conditions. As in previous studies, the statistical significance of choice probabilities was determined relative to chance. We shuffled behavioral bout timings for each of the two compared conditions and computed the choice probability for this shuffled data. Shuffling was repeated 100 times for each of the two behaviors, from which we calculated the mean and s.d. (*σ*) of the ‘shuffled’ choice probabilities.

As significant, we considered any observed choice probabilities >2*σ* above the shuffled mean and imposed an additional choice probability threshold > 0.7 as previously described^58^. The colored bars indicate the neurons that show a strong and statistically significant choice probability, and grey bars indicate cells for which the choice probability was either activated < 2 σ above (not responsive) the shuffled mean or was considered which choice probability not significantly higher than chance or choice probability ≤ 0.7 for that neuron.

### Dimensionality reduction for visualizing intruder sex

Low-dimensional representations for visualizing changing ensemble dynamics over time were constructed using partial least squares (PLS) regression (MATLAB). For PLS, all traces were concatenated and regressed against a 1 × *T* vector with entries valued at −1 (if a male intruder was present), 1 (if there was a female intruder), or 0 (otherwise).

### Decoding intruder sex from neural data

We constructed a frame-wise linear SVM decoders (as described previously^58,59^) to distinguish intruder sex. Training data was constructed from the set of *N* × 1 (*N* = neurons) population activity vectors from all frames occurring during social interaction in each mouse. Equal numbers of frames of male and female interaction were used during decoder training to ensure chance decoder performance of 50%. Shuffled decoder data were generated by the training the decoder on the same neural data but with behavior labels randomly assigned to each behavior bout (n=5 control and n=7 *expreimental mice*). This training data, along with intruder sex labels, was then used to train a linear SVM decoder. Accuracy was evaluated using a stratified fivefold cross-validator. Decoding was repeated 100 times, with decoder performance reported as the mean accuracy per imaged animal. For significance testing, the mean accuracy of the decoder trained on shuffled data (repeated 500 times per imaged animal) was computed to compare against the decoder accuracy trained on actual data.

### Decoding behavior from neural activity

We constructed frame-wise linear SVM decoders (as described previously^57,58^) to discriminate male directed sniffing and attack from imaged control and experimental VMHvl^Esr1^ units. Briefly manual annotations of sniffing behavior and attack behavior for each intruder male mouse were used to provide training labels of behavior type in control and experimental mice. Bar graphs of decoder accuracy (Figure S5E) were generated to discriminate sniffing and attack from imaged activity on individual frames of a behavior (sampled at 15 Hz). Equal numbers of sniff and attack frames (frame-wise decoder) were used during decoder training, to ensure chance decoder performance of 50%. ‘Shuffled’ decoder data were generated by training the decoder on the same neural data, but with sniff and attack behavior annotations randomly assigned to each behavior bout.

Decoding was repeated 20 times for each intruder and each imaged mouse, and decoder performance was reported as the average accuracy across imaged mice for control and experimental mice. For significance testing, the mean accuracy of the decoder trained on shuffled data was computed across mice, in each condition, and shuffling was repeated 1,000 times. Significance was determined across imaged mice using the Mann–Whitney U test between the mean accuracy of the decoders trained on real versus shuffled data.

### Statistical analysis

Data were processed and analyzed using Python, MATLAB and GraphPad (GraphPad PRISM v.9). Data were analyzed using two-tailed, nested non-parametric tests. Wilcoxon signed-rank test (paired, non-parametric Mann–Whitney *U*-test) was used for binary paired samples. Kolmogorov–Smirnov test was used for non-paired samples plotted as ECDF graphs. N.s. *P* > 0.05, \**P* < 0.05, \*\**P* < 0.01, \*\*\**P* < 0.001, \*\*\*\**P* < 0.0001.

### Dynamical system models of neural data

As previously described in published work^20^, we modeled neural activity using recurrent switching linear dynamical systems (rSLDS). Briefly, rSLDS is a generative model that breaks down non-linear time series data into sequences of linear dynamical modes. The model relates three sets of variables: a set of discrete states (z), a set of continuous latent factors (x) that captures the low-dimensional nature of neural activity, and the activity of recorded neurons (y). The model also allows for external inputs (u), which consist of extracted pose features, including the distance between animals and the facing angle between the resident and intruder mouse. For how the model is formulated, see Nair, et al., 2023^20^. Model accuracy is evaluated using a forward simulation metric as described in Nair et al., 2023^20^. Briefly: given the observed neural activity at time *t*, we predict the trajectory of the population activity vector over an ensuing short time interval Δ*t* using the model, then compute the mean squared error (MSE) between that trajectory and the observed data at time *t+* Δ*t*. This MSE is calculated across all dimensions of the latent space and repeated for all times *t*. This error metric is normalized to a 0-1 range in each animal across the whole recording and is computed across cross-validation folds to obtain a bounded measure of model performance.

Code used to fit rSLDS on neural data is available in the SSM package: (https://github.com/lindermanlab/ssm)

Code to generate flow fields and energy landscapes from fit dynamical systems is available at (https://github.com/DJALab/VMHvl_MPOA_dynamics)

### Visualization of attractor dynamics as 3D landscape

Conversion of the flow-fields obtained from rSLDS into a 3D landscape for visualization by calculating the dynamic velocity at each point in neural state space and using it as the height of a 3D landscape. Dynamic velocity was calculated as previously reported in *Nair et al., 2022*.

### Estimation of time constants & calculation of line attractor score

We estimated the time constant of each mode of linear dynamical systems using eigenvalues λ_*a*_ of the dynamics matrix of that system as: 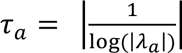 as derived by Maheswaranathan et al., 2019^121^. We used a line attractor score computed as 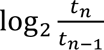 where *t*_*n*_ is the largest time constant of the dynamics matrix of a dynamical system and*t*_*n*–1_ is the second largest time constant. In the case of point attractors, the line attractor score is zero due to the similar magnitudes of the first two largest time constants, and it is greater than one for systems that possess a line attractor.

