## Supplementary figures and images for "Neuropeptide Signaling is Required to Implement a Line Attractor Encoding a Persistent Internal Behavioral State"

### Supplemental figures

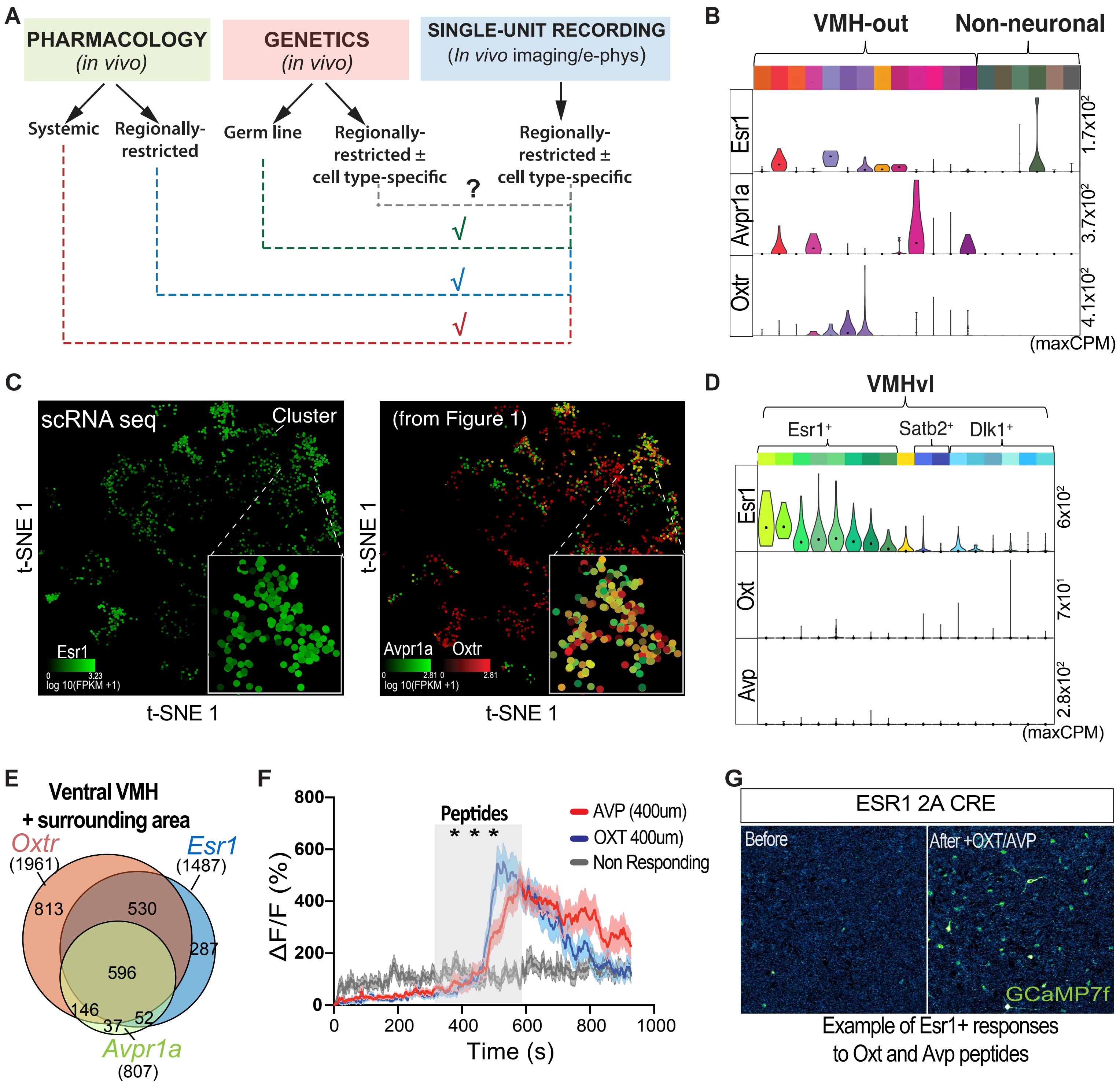

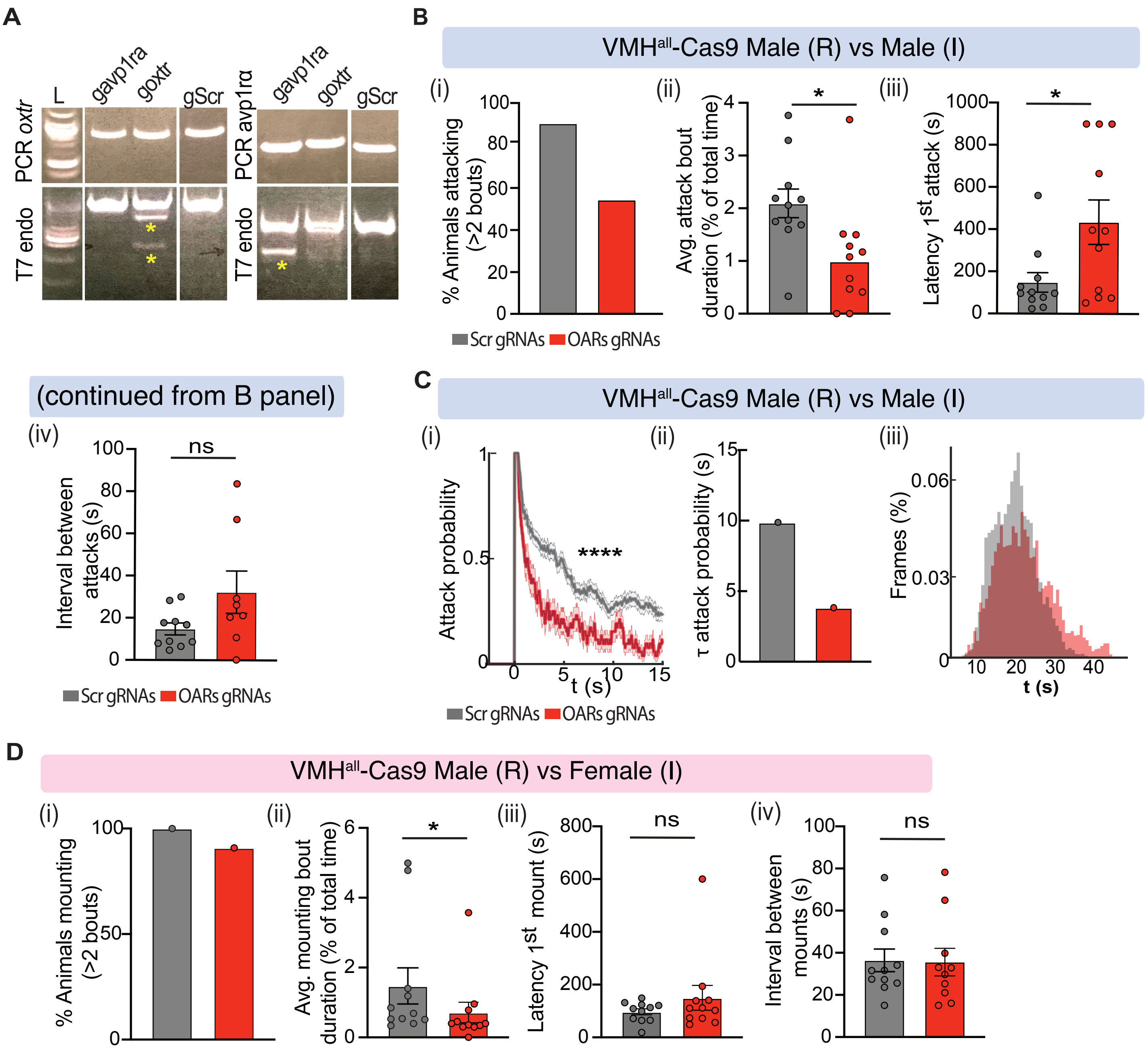

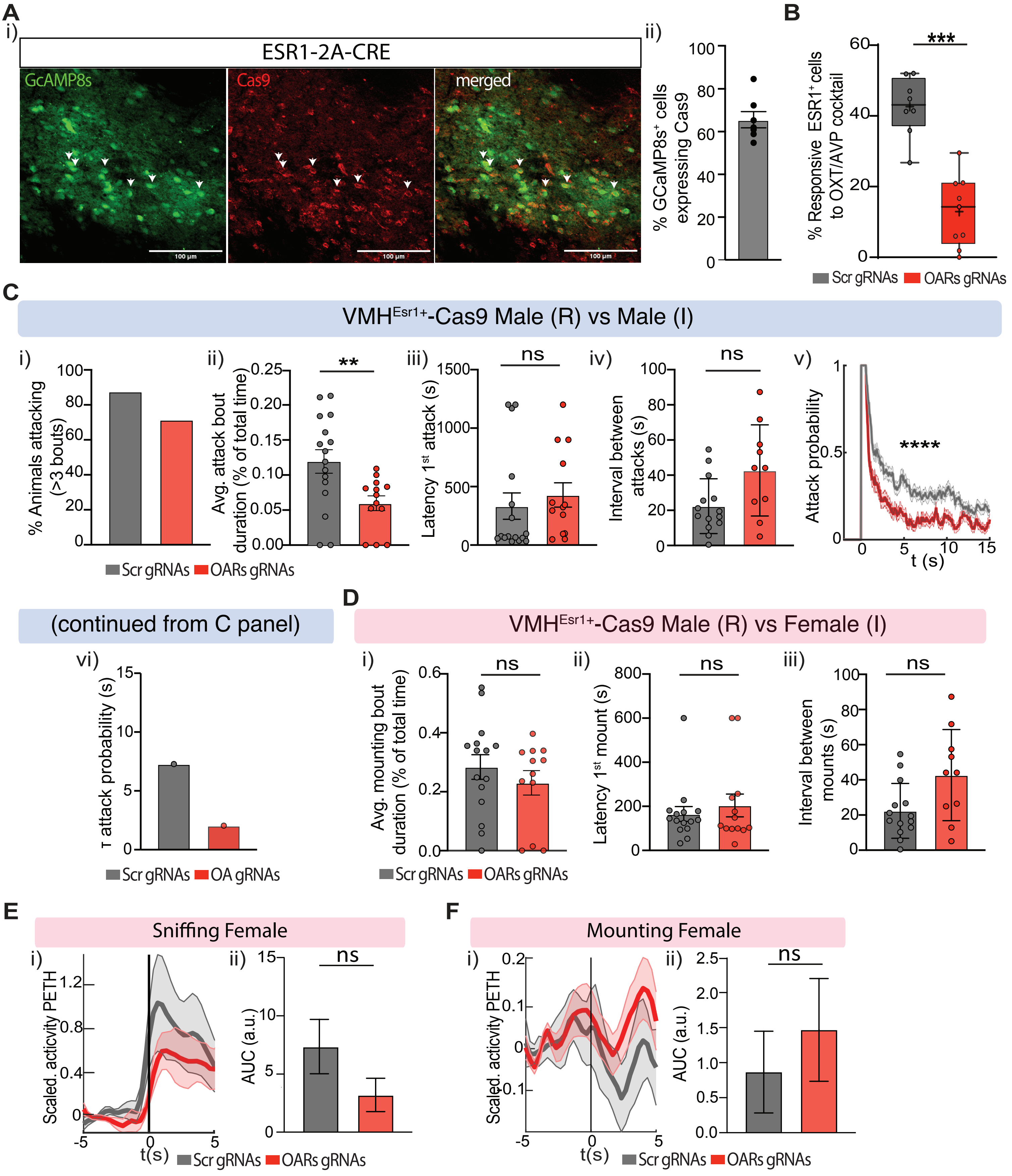

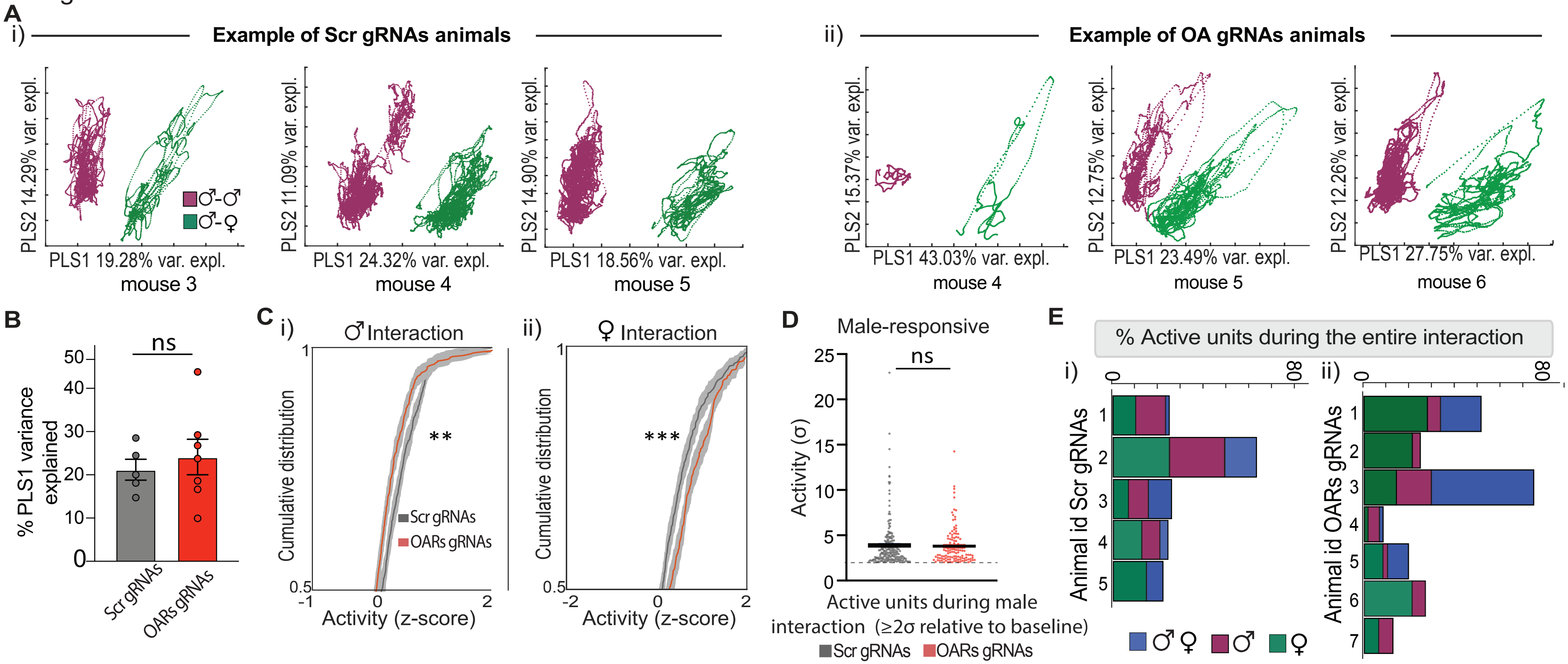

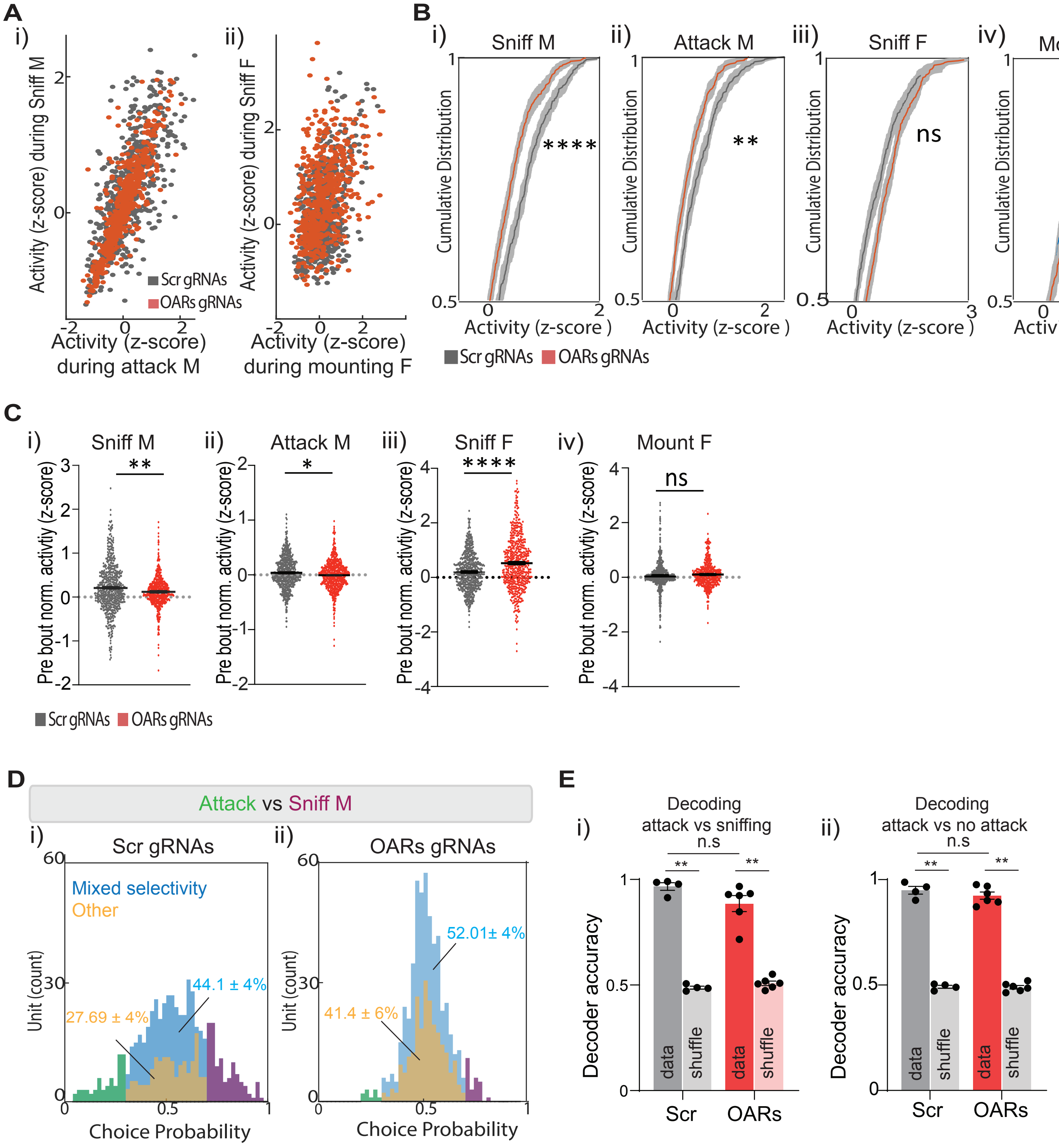

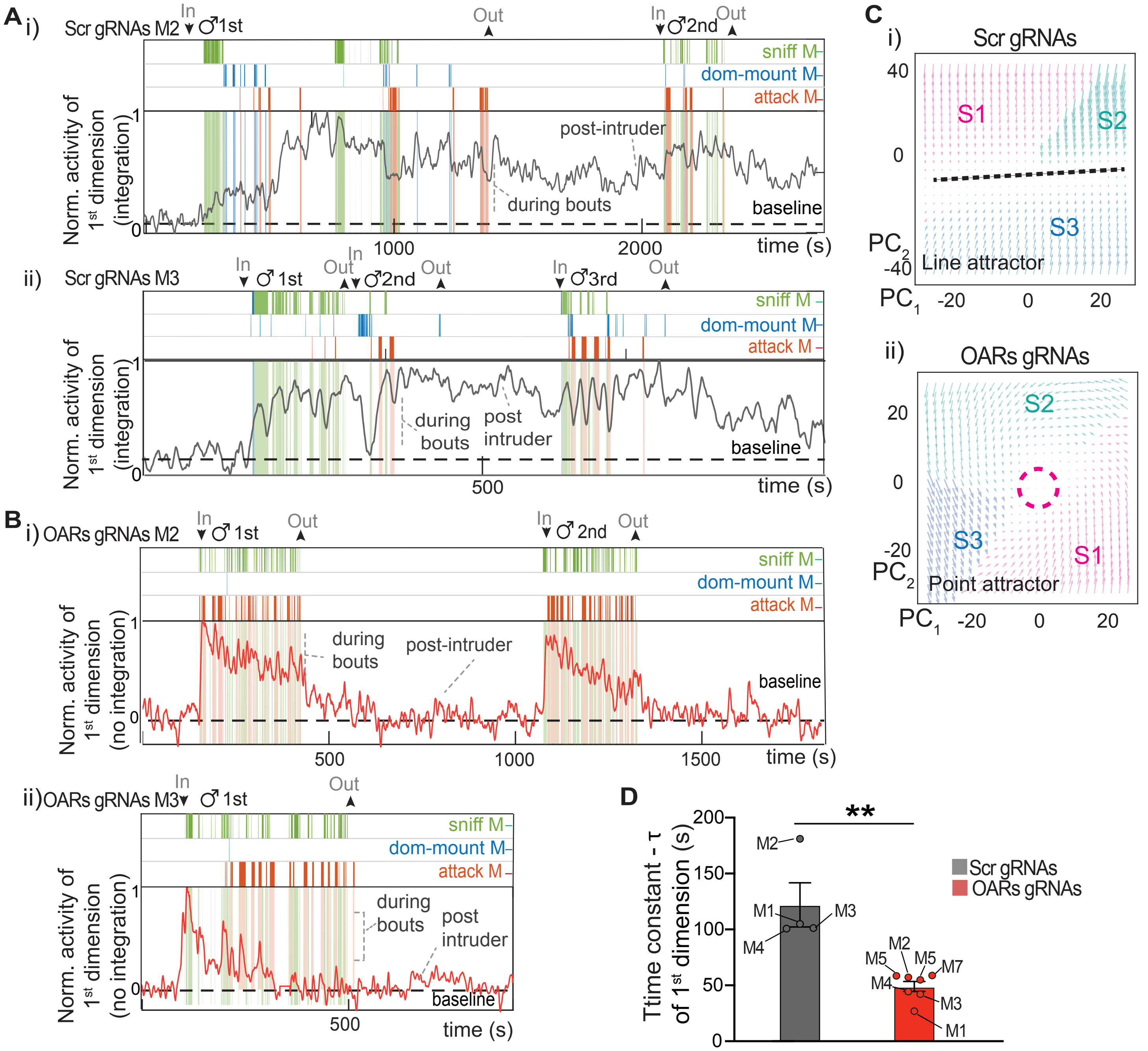

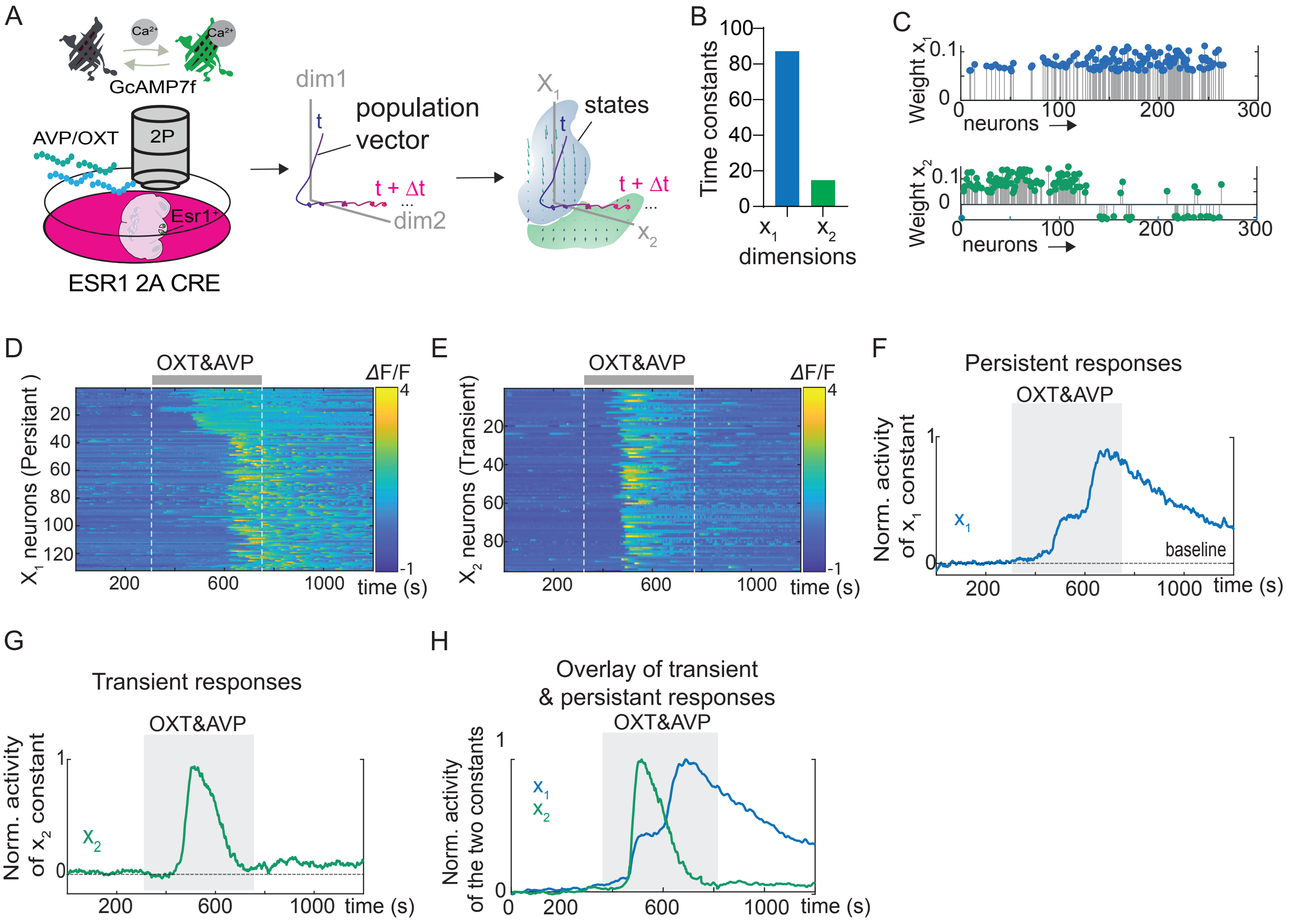
